# Effect of homotypic *vs*. heterotypic interactions on the cellular uptake of extracellular vesicles

**DOI:** 10.1101/2023.10.23.563628

**Authors:** Jhanvi R. Jhaveri, Purva Khare, Paromita Paul Pinky, Yashika S. Kamte, Manisha N. Chandwani, Jadranka Milosevic, Nevil Abraham, Kandarp M. Dave, Si-yang Zheng, Lauren O’Donnell, Devika S Manickam

## Abstract

Extracellular vehicles (**EVs**) are an emerging class of drug carriers and are primarily reported to be internalized into recipient cells via a combination of endocytic routes such as clathrin-mediated, caveolae-mediated and macropinocytosis pathways. In this work, (*1*) we investigated potential effects of homotypic *vs*. heterotypic interactions by studying the cellular uptake of homologous EVs (EV donor cells and recipient cells of the same type) *vs*. heterologous EVs (EV donor cells and recipient cells of different types) and (*2*) determined the route of EV internalization into low pinocytic/hard-to-deliver cell models such as brain endothelial cells (**BECs**). We used BECs and macrophages as low-pinocytic and phagocytic cell models, respectively, to study the effect of homotypic *vs*. heterotypic interactions on EV uptake in the recipient cells. Homotypic interactions led to a greater extent of uptake into the recipient BECs compared to heterotypic interactions. However, we did not see a complete reduction in EV uptake into recipient BECs when endocytic pathways were blocked using pharmacological inhibitors. Our results suggest that EVs primarily use membrane fusion to enter low-pinocytic recipient BECs instead of relying on endocytosis. *Lipophilic PKH67 dye-labeled EVs* but not intravesicular esterase-activated calcein ester-labeled EVs severely reduced particle uptake into BECs while phagocytic macrophages internalized both types of EV-labeled particles to comparable extents. Our results also highlight the importance of carefully choosing labeling dye chemistry to study EV uptake, especially in the case of low pinocytic cells such as BECs.

**Graphical abstract:** 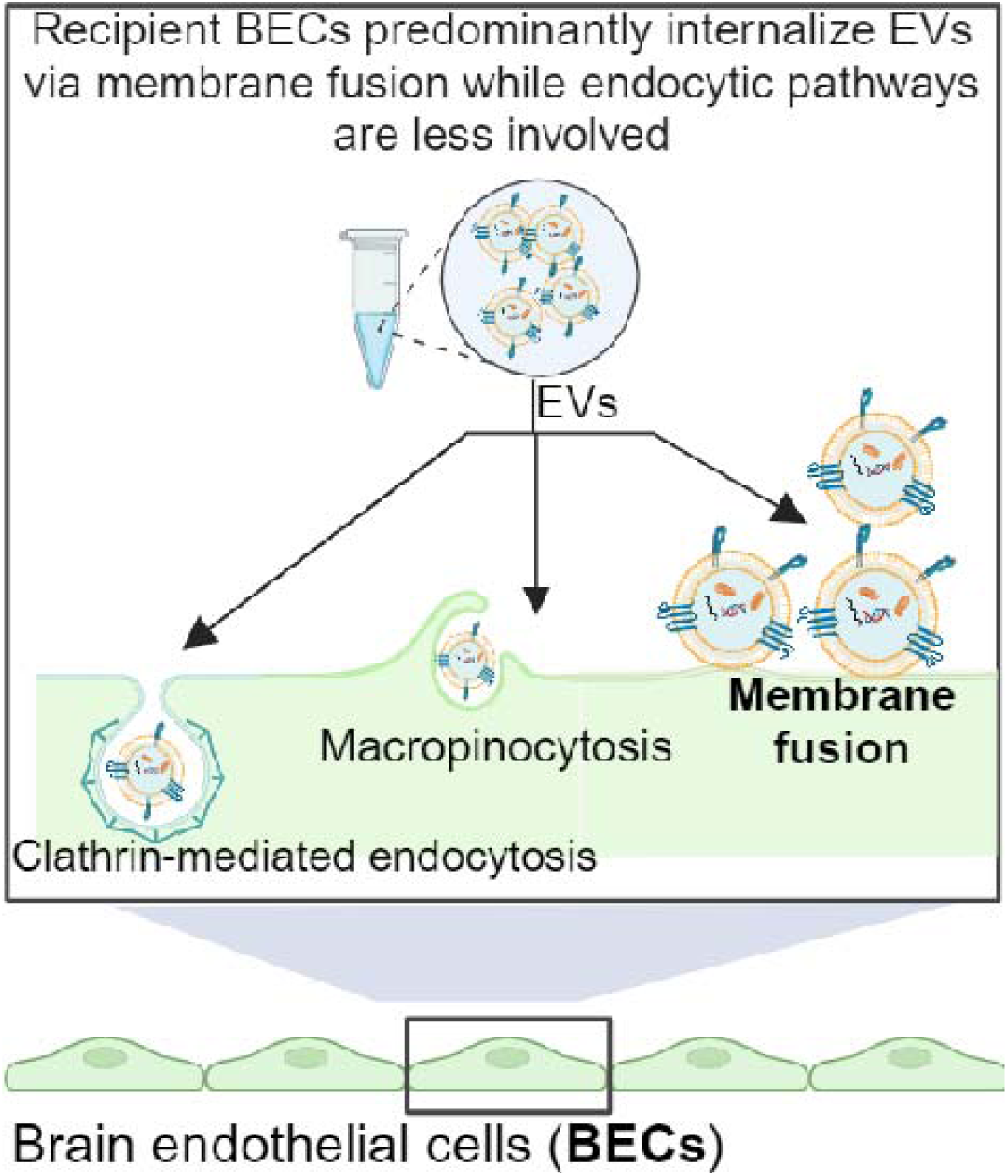

## 1. Introduction

Extracellular vesicles (**EVs**) transfer cellular cargo via the transfer of proteins, nucleic acids, lipids and other vesicular contents between donor and recipient cells [1–5]. The most widely studied EVs are of two subtypes vary in EV particle diameter in addition to the different biogenesis pathways involved in their secretion into extracellular spaces: the 100-1000 nm medium-to-large EVs (**m/lEVs**) that are sometimes referred to as microvesicles and the 50-200 nm small EVs (**sEVs**), commonly referred to as exosomes [6]. The biogenesis of sEVs involves inward budding of the endosomal membrane into intraluminal vesicles and are released in the extracellular environment by fusion of multivesicular endosomes with the cell membrane. m/lEVs are formed by the outward budding of the cell membrane [4, 7]. The well-documented role of EVs in intercellular communication propelled the development of EVs as carriers for drug delivery [6, 8, 9]. Their biological/natural origin and potential for lower immunogenicity may allow improved drug delivery compared to synthetic nanoparticles.

EVs retain membrane signatures (lipids and proteins) reminiscent of their donor/parent cells that may allow increased uptake into recipient cells of the same type [10, 11]. For clarity, we define the similarity of the EV donor/recipient cell pairs as “homotypic” interactions whereas the use of different EV donor and recipient cell pairs are referred to as “heterotypic” interactions. Studying the effects of homotypic/heterotypic interactions on EV uptake into the recipient cells is important to guide the proper selection of EV donor cells and maximize effects of EV uptake and delivery. Our laboratory has previously demonstrated that m/lEVs derived from brain endothelial cells (**BECs**) increase mitochondrial function in the recipient BECs and also result in reduced brain tissue infarct volumes in a mouse model of transient ischemic stroke [12–15]. We have demonstrated that BEC-derived EVs (**BEC-EVs**) show a greater extent of fluorescent mitochondria signals in the recipient BECs as opposed to EVs derived from a macrophage cell line. Concurrent with the increased fluorescent mitochondria signals, recipient BECs treated with BEC-EVs showed a greater extent of ATP increase compared to recipient BECs treated with macrophage-derived EVs [15]. These results seem to suggest that the interactions between homotypic EVs and recipient cells (recipient BECs treated with BEC-EVs) *vs*. interactions between heterotypic EVs and the recipient cells (recipient BECs treated with macrophage-derived EVs) likely play a role in the functional EV responses in the recipient cells. In this work, we sought to [1] determine if there are any potential differences in the cellular uptake of homotypic *vs*. heterotypic EVs into recipient BECs and [2] determine the potential mechanism of EV cellular uptake into low pinocytic cells such as BECs.

Recipient cells are mostly known to internalize EVs using various endocytic routes including clathrin-dependent as well as clathrin-independent pathways such as macropinocytosis and phagocytosis [4, 16–18] but only a couple of reports have indicated the role of membrane fusion between EV and cell membranes [19, 20]. Noteworthy, published reports use EVs from various cell sources and have determined entry/uptake routes in different recipient cell models [5, 16, 21–23]. For instance, Costa Verdera *et al*. showed that the A431 human epidermoid carcinoma-derived EVs enter the recipient human cervical carcinoma HeLa cells via clathrin-independent endocytosis and macropinocytosis routes [24]. Ilahibaks *et al*. showed clathrin-dependent endocytosis and macropinocytosis-mediated delivery of human embryonic kidney 293FT (HEK293FT)-derived EVs [25] into T47D stoplight reporter cells. The uptake route of BEC-derived EVs into recipient BECs is of specific interest to us as we continue to develop these EVs as a unique class of mitochondria carriers [12–15, 26].

In this work, (1) we studied the cellular uptake of EVs labeled using either a lipophilic PKH67 dye or calcein acetoxymethyl (**calcein AM**) ester dye activated by intravesicular esterases in membrane-intact EVs to determine the potential effect of labeling dye chemistry on the extent of cellular uptake and (2) we blocked three key routes of endocytosis: clathrin-mediated; caveolae-mediated and macropinocytosis, using their respective pharmacological inhibitors to study the uptake mechanism of homotypic *vs.* heterotypic EVs into recipient BECs or macrophages. Our results suggested that blocking different endocytosis pathways modestly decreased EV uptake into the recipient BECs, however, it did not completely block the EV uptake, suggesting the involvement of energy-independent pathways like membrane fusion. A R18 self-quenching dye-based fusion assay demonstrated that the early phase (first 30-minutes) of BEC-EV uptake into recipient BECs occurs via membrane fusion. PKH67-but *not* calcein-dye labeled EVs showed reduced uptake into BECs—underscoring the importance of choosing appropriate dye chemistry to study cellular uptake in low pinocytic cell models.

## 2. Materials and methods

### Key reagents

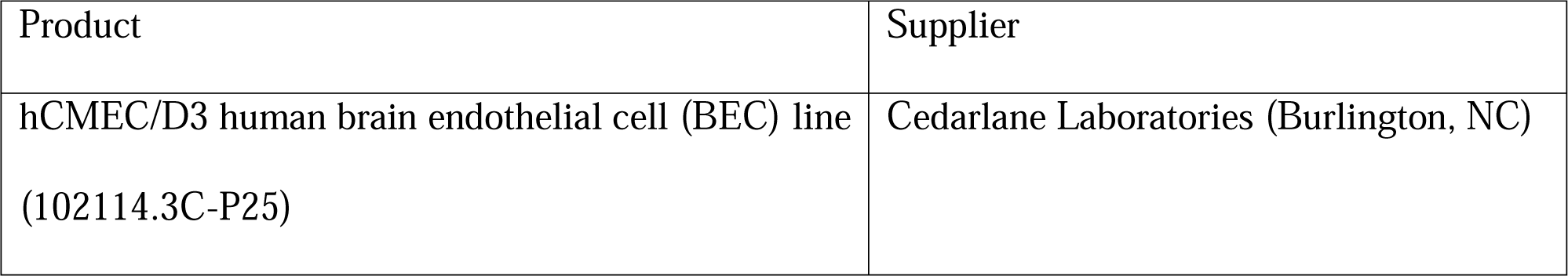
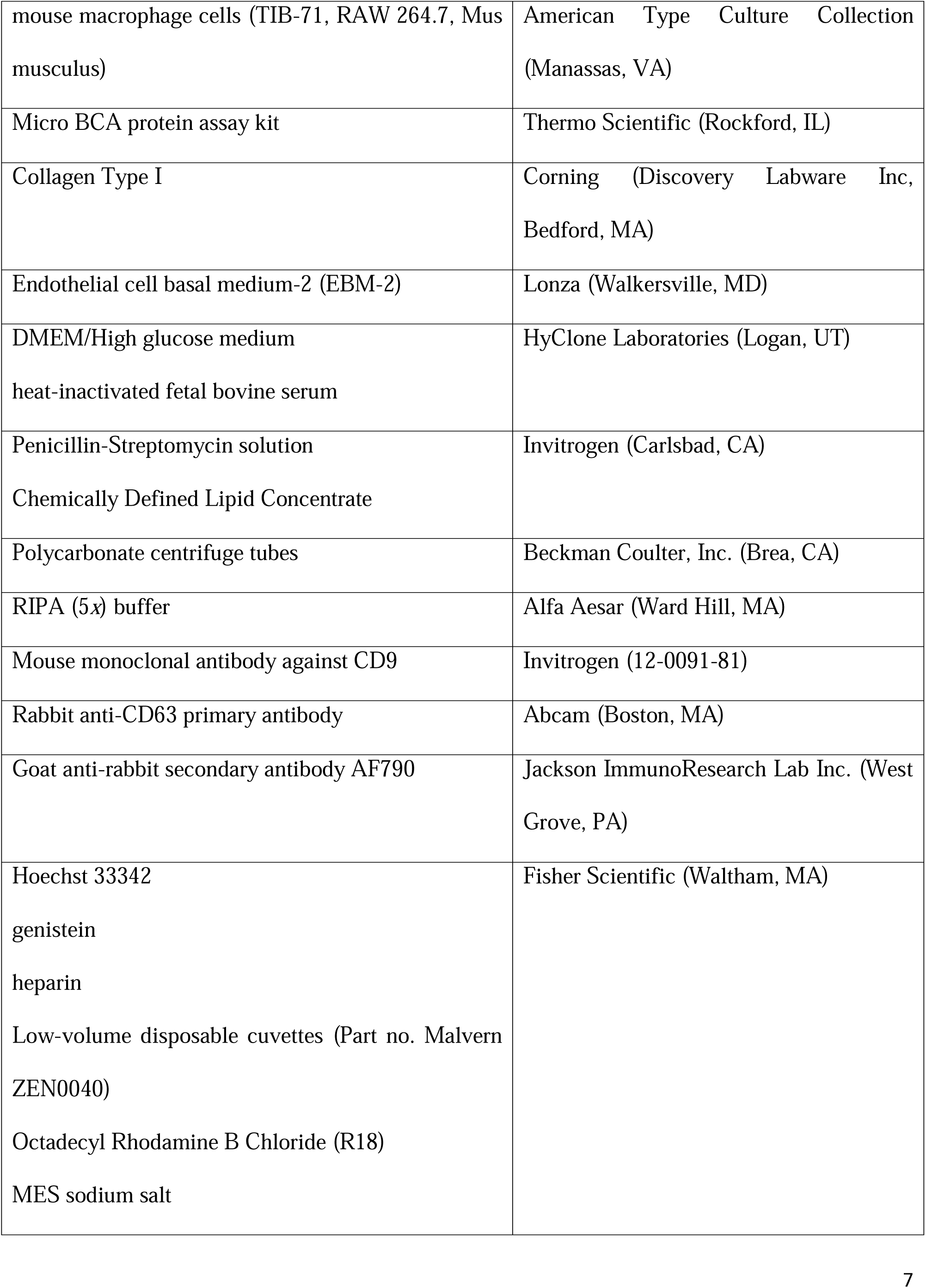
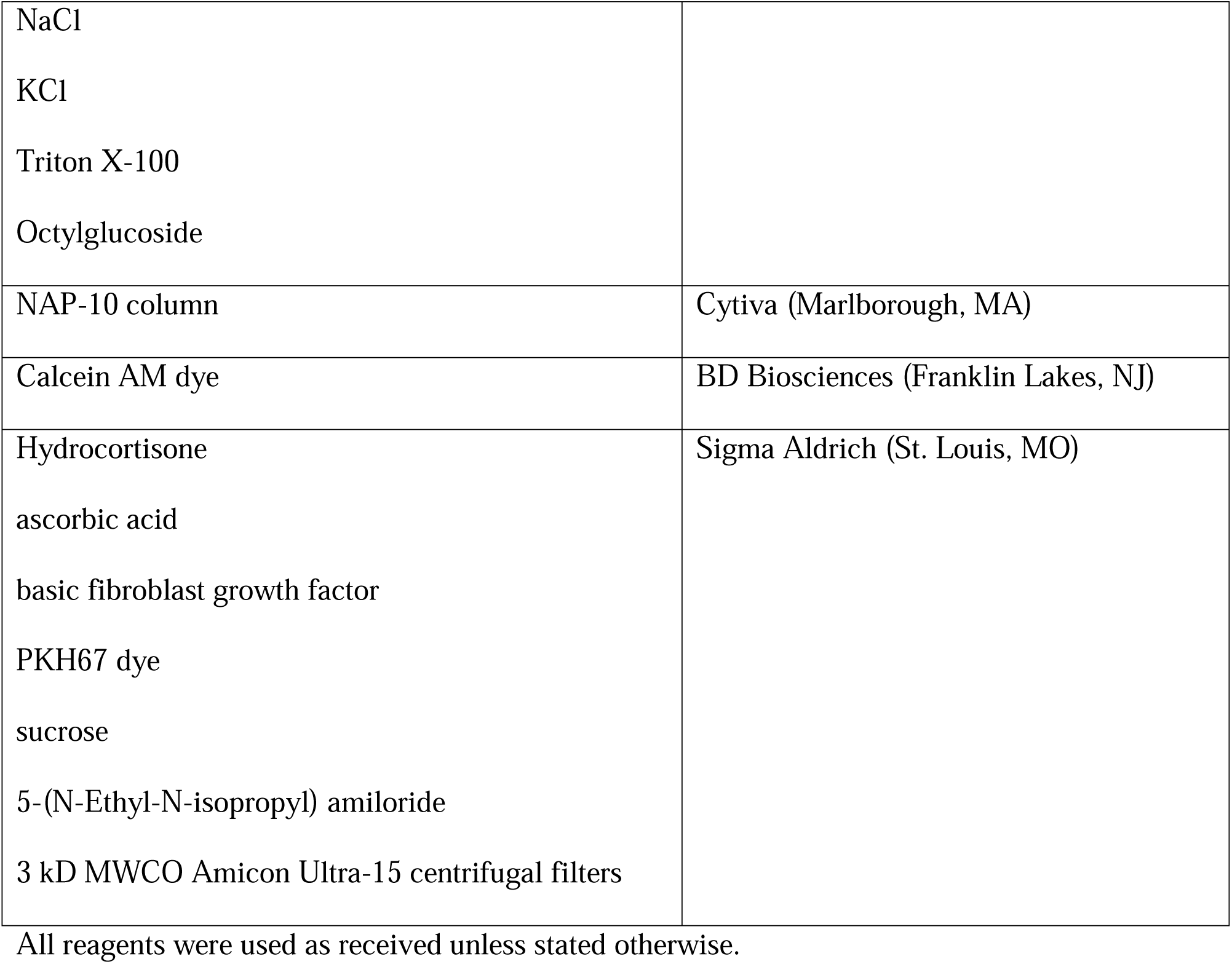

| Product | Supplier |
| --- | --- |
| hCMEC/D3 human brain endothelial cell (BEC) line<br>(102114.3C-P25) | Cedarlane Laboratories (Burlington, NC) |
| mouse macrophage cells (TIB-71, RAW 264.7, Mus musculus) | American Type Culture Collection (Manassas, VA) |
| Micro BCA protein assay kit | Thermo Scientific (Rockford, IL) |
| Collagen Type I | Corning (Discovery Labware Inc, Bedford, MA) |
| Endothelial cell basal medium-2 (EBM-2) | Lonza (Walkersville, MD) |
| DMEM/High glucose medium<br>heat-inactivated fetal bovine serum | HyClone Laboratories (Logan, UT) |
| Penicillin-Streptomycin solution<br>Chemically Defined Lipid Concentrate | Invitrogen (Carlsbad, CA) |
| Polycarbonate centrifuge tubes | Beckman Coulter, Inc. (Brea, CA) |
| RIPA (5x) buffer | Alfa Aesar (Ward Hill, MA) |
| Mouse monoclonal antibody against CD9 | Invitrogen (12-0091-81) |
| Rabbit anti-CD63 primary antibody | Abcam (Boston, MA) |
| Goat anti-rabbit secondary antibody AF790 | Jackson ImmunoResearch Lab Inc. (West Grove, PA) |
| Hoechst 33342<br>genistein<br>heparin<br>Low-volume disposable cuvettes (Part no. Malvern ZEN0040)<br>Octadecyl Rhodamine B Chloride (R18)<br>MES sodium salt | Fisher Scientific (Waltham, MA) |
| NaCl |  |
| KCl |  |
| Triton X-100 |  |
| Octylglucoside |  |
| NAP-10 column | Cytiva (Marlborough, MA) |
| Calcein AM dye | BD Biosciences (Franklin Lakes, NJ) |
| Hydrocortisone | Sigma Aldrich (St. Louis, MO) |
| ascorbic acid |  |
| basic fibroblast growth factor |  |
| PKH67 dye |  |
| sucrose |  |
| 5-(N-Ethyl-N-isopropyl) amiloride |  |
| 3 kD MWCO Amicon Ultra-15 centrifugal filters |  |

All reagents were used as received unless stated otherwise.

### 2.1 Cell culture

hCMEC/D3 BECs were maintained in tissue culture flasks or multi-well plates pre-coated with 0.15 mg/mL rat collagen I in a humidified 5% CO_2_ incubator at 37 ± 0.5°C (Isotemp, Thermo Fisher Scientific). The cells were cultured in complete growth medium comprised of endothelial cell basal medium (EBM-2) supplemented with fetal bovine serum (5% FBS), penicillin (100 units/mL)-streptomycin (100 μg/mL) mixture, hydrocortisone (1.4 µM), ascorbic acid (5 µg/mL), Chemically Defined Lipid Concentrate (1%), basic fibroblast growth factor (1 ng/mL) and 10 mM HEPES (pH 7.4). The complete growth medium was renewed every other day until the cells formed confluent monolayers. RAW 264.7 cells were maintained in tissue culture flasks or multi-well plates in a humidified 5% CO_2_ incubator at 37 ± 0.5°C (Isotemp, Thermo Fisher Scientific). The cells were grown in DMEM/high glucose medium supplemented with fetal bovine serum (10% FBS). Prior to passaging, the cells were washed using 1*x* phosphate buffer saline (**PBS**) and were detached using 1*x* TrypLE Express Enzyme (Gibco, Denmark). hCMEC/D3 BECs maintained between P25 and P35 and RAW 264.7 macrophages maintained between P7 and P25 and were used in all experiments.

### 2.2 Isolation of sEVs and m/lEVs

<u>Both large (m/lEVs) and small (sEVs) EV fractions are collectively referred to as EVs</u> wherever applicable. We used a differential ultracentrifugation method to isolate m/lEVs and sEVs from the EV-conditioned supernatants of BECs and macrophages [12, 15, 27, 28]. Briefly, four tissue culture flasks of 182 cm^2^ growth area (T-175) containing BECs/macrophages were washed with pre-warmed 1*x* PBS and incubated with serum-free medium (complete growth medium lacking FBS and penicillin-streptomycin) for 48 h in a humidified 5% CO_2_ incubator at 37 ± 0.5°C. Post-incubation, a total of 80 mL EV-conditioned medium was collected in polypropylene centrifuge tubes and initially centrifuged at 300 ×g for 11 minutes at 4°C to pellet down dead cells followed by subsequent centrifugation of the supernatant at 2000 ×g for 22 min at 4°C to pellet down apoptotic bodies and cell debris using a Sorvall ST 8R centrifuge (ThermoFisher Scientific, Osterode am Harz, Germany). The supernatant was then transferred into polycarbonate tubes and centrifuged at 20,000 ×g for 45 min at 4°C to pellet down m/lEVs using an Optima XE-90 ultracentrifuge fitted with a 50.2 Ti rotor (Beckman Coulter, Indianapolis, IN). Following that, the supernatant was filtered through a 0.22 µm PES membrane syringe filter, and the flow-through was centrifuged at 120,000 ×g for 70 min at 4°C to collect sEVs. Finally, m/lEV and sEV pellets were suspended in 1.2 mL 1*x* PBS, respectively. Furthermore, EVs were suspended in 1*x* PBS for particle diameter measurements and *in vitro* experiments or DI water for zeta potential measurements. m/lEV and sEV samples were stored at −20°C until further use. Pierce MicroBCA assay was used to quantify the total protein content in m/lEVs and sEVs. Briefly, m/lEVs and sEVs were lysed using 1*x* RIPA buffer supplemented with 3 μg/mL aprotinin in a 1:15 v/v ratio. Next, a 150 μL volume of m/lEV or sEV lysates or BSA standards (0.5–200 μg/mL) was pipetted into a 96-well plate and an equal volume of the MicroBCA working reagent (reagent A: reagent B: reagent C at 25:24:1 volume ratio) was added to each well. This mixture was incubated at 37°C for 2 h and the absorbance was measured at 562 nm using a SYNERGY HTX multi-mode reader (BioTek Instruments Inc., Winooski, VT).

### 2.3 Dynamic light scattering (DLS) and Nanoparticle tracking analysis (NTA)

Particle diameter and zeta potential of EVs were measured using DLS on a Malvern Zetasizer Pro (Malvern Panalytical Inc., Westborough, PA). EV particle concentrations were determined on a multiple-laser ZetaView f-NTA Nanoparticle Tracking Analyzer (Particle Metrix Inc., Mebane, NC). For DLS, EVs were prepared at 0.1 mg protein/mL suspended in 1*x* PBS and DI water at pH 7.4 for particle size and zeta potential measurements, respectively. Average particle diameter, dispersity index, and zeta potential values were reported as mean ± standard deviation. For NTA, stock samples of m/lEVs and sEVs were diluted in 1*x* PBS before analysis. Five 60 s videos were acquired at 520 nm laser wavelength for concentration measurements. Data were reported as mean ± standard deviation.

### 2.4 Multispectral imaging flow cytometry

Characteristic EV markers were identified by performing flow-cytometric multispectral imaging of BEC and macrophage-derived sEVs and m/lEVs using an ImageStreamX MKII flow cytometer. EVs (1x 10^8^ particles/mL) were labeled with CD9 (1 µg) antibody. CD9 stock tube was centrifuged at 16,000 xg for 10 minutes at 4°C to remove any aggregates. EVs were incubated with the antibody for 60 minutes at room temperature in the dark. The acquisition of EVs using ImageStreamX was performed with fluidics set at low speed, magnification at 60×, core size 7 µm, and the “Hide Beads” option unchecked before every acquisition to visualize speed beads using INSPIRE software. ImageStreamX was equipped with the following lasers: 405 nm, 488 nm, 561 nm, and 642 nm, and the 488 nm laser was active during the experiment. Ch01 and Ch09 were set to brightfield (BF), allowing spatial coordination among cameras. Channel 06 was set to side-scatter (SSC) to gate total EVs out of all events (including speed beads), and fluorescence channel (Ch03) was used to detect the antibody signals.

### 2.5 Western Blotting

Characteristic EV markers were detected by western blotting using our previously reported protocol [12, 13, 15, 27]. Briefly, cell and EV lysates comprising of 20 and 70 μg total protein, respectively were mixed with 4× Laemmli buffer and distilled water. This mixture was heated at 95 °C for 5 min using a heating block (Thermo Scientific). The samples and the premixed molecular weight markers (ladder, 250 kD-10 kD) were separated on a 4–10% gradient sodium dodecyl sulfate-polyacrylamide gel at 120 V for 90 min using a PowerPac Basic setup (Bio-Rad Laboratories, Inc.). Next, the proteins were transferred to a 0.45-μm nitrocellulose membrane using a transfer assembly (Fisher Scientific) at 75 V and 300 mA for 90 min. Next, the membrane was washed with the 0.1%-Tween 20 Tris buffered-saline (T-TBS) and blocked using Odyssey blocking solution (Odyssey blocking buffer: T-TBS, 1:1) solution for an hour. The membrane was incubated overnight at 4 °C with rabbit anti-CD63 primary antibody (1 mg/mL) at (1/300 dilution in Odyssey blocking solution). The membrane was again washed with T-TBS and incubated with anti-rabbit AF790 (1:5000 dilution in Odyssey blocking solution) secondary antibody at room temperature for an hour. The washed membrane was scanned at 700 and 800-nm near-infrared channels using an Odyssey M imager (LI-COR Inc., Lincoln, NE).

### 2.6 Labeling of EVs using calcein AM or PKH67

EVs suspended in 1*x* PBS were labeled with 10 μM calcein AM for 20 min at room temperature in dark or with 5 μM PKH67 according to the manufacturer’s instructions [24]. To remove the excess/free PKH67 dye post-labeling the EVs, pooled EV fractions were concentrated at 4°C using 3 kD MWCO Amicon Ultra-15 Centrifugal Filter Units and the calcein AM-labeled EVs were used as such.

### 2.7 Uptake of calcein AM- or PKH67-labeled EVs into recipient BECs or macrophages using flow cytometry

EVs suspended in 1*x* PBS were labeled with 10 μM calcein AM or with 5 μM PKH67 as described in section 2.5. BECs/RAW macrophages were cultured in 24-well plates at 90,000 cells/well in their respective complete growth medium. Post-confluency, the cells were treated with either calcein AM- or PKH67-labeled EVs at 5, 25, and 50 µg EV protein/well in complete growth medium for 4 h in a humidified incubator. Untreated cells were used as gating control. Post-treatment, the cells were washed with 1*x* PBS and an acid buffer (0.5 M NaCl and 0.2 M CH_3_COOH) to remove any non-internalized EVs bound to cell membranes. The cells were then dissociated using TrypLE Express, suspended in 3% FBS in 1*x* PBS, and collected in eppendorf tubes. For each sample, a 50 µL aliquot of the cell suspension was analyzed using an Attune NxT flow cytometer and at least > 15,000 events were recorded in FSC *vs*. SSC plots. Calcein AM or PKH67 fluorescence intensity was detected at 488/10 nm and percentage signal intensities were recorded for data analysis.

### 2.8 Uptake of calcein AM- *vs.* PKH67-labeled-EVs into recipient macrophages using fluorescence microscopy

EVs suspended in 1*x* PBS were labeled with calcein AM or PKH67 as discussed in section 2.5. RAW macrophages were cultured in 24-well plates in DMEM/high glucose medium supplemented with 10% FBS. Post-confluency, the cells were treated with calcein AM or PKH67-labeled EVs at 1, 5, 25 or 50 µg protein/well in complete medium for 4 h in a humidified incubator. Prior to observation, the treatment mixture was removed, and the cells were washed with 1*x* PBS and acid buffer followed by addition of 1 mL phenol-red free complete growth medium and the nuclei was stained with 10 µg/mL Hoechst dye for 10 minutes. Untreated cells were used as the negative control. Cells were observed under an Olympus IX 73 epifluorescent inverted microscope (Olympus, Pittsburgh, PA) using GFP channel to detect calcein AM and PKH67 signals and DAPI channel to detect Hoechst signals at 20*x* magnification. Images were obtained and quantified using CellSens Dimension software (Olympus, USA).

### 2.9 Uptake of calcein AM-labeled EVs into recipient BECs using z-stack fluorescence microscopy

EVs suspended in 1*x* PBS were labeled with calcein AM as discussed in section 2.5. BECs were cultured in 24-well plates. Post-confluency, the cells were treated with calcein AM-labeled EVs at 25 µg protein/well in complete medium for 24 h in a humidified incubator. Prior to observation, the cells were washed and stained with Hoechst dye as described in section 2.7. Untreated cells were used as the negative control. Cells were observed under an Olympus IX 73 epifluorescent inverted microscope (Olympus, Pittsburgh, PA) using GFP channel to detect calcein AM signals and DAPI channel to detect Hoechst signals at 20x magnification. Z-stack images were used to visualize the intracellular presence of EVs in the recipient BECs. Images were obtained using CellSens Dimension software (Olympus, USA).

### 2.10 Uptake of calcein AM-labeled-EVs into recipient BECs and macrophages treated with endocytosis inhibitors

BECs and macrophages were cultured in 24-well plates at 90,000 cells/well in their complete growth medium, respectively. Cells were preincubated for 30 minutes with either genistein (200 µM), sucrose (250 mM) or 5-(N-Ethyl-N-isopropyl) amiloride (**EIPA**, 100 µM) (endocytosis inhibitors) before EV treatment. This was followed by a co-treatment with calcein AM-labeled EVs and fresh endocytosis inhibitors for 4 h at either 25 µg or 50 µg EV protein/well, for recipient BECs and macrophages, respectively. The cells were harvested for flow cytometry experiment as described in section 2.6. Endocytosis inhibitors were present throughout the experiment. The % EV uptake and median fluorescence intensity (**MFI**) were recorded for data analysis.

### 2.11 Effect of heparin on the uptake of calcein AM-labeled EVs into recipient BECs

BECs were cultured in 24-well plates at 100,000 cells/well in complete growth medium. Cells were preincubated with heparin (100 µg/mL) or 1*x* PBS (control) before EV treatment for 30 minutes. This was followed by a co-treatment with calcein AM-labeled EVs (25 µg EV protein/well) and fresh heparin or 1*x* PBS for either 30 minutes or 4 h. The cells were harvested and analyzed using flow cytometry experiment as described in section 2.6 and 2.9.

### 2.12 Membrane fusion assay

EV fusion with the recipient BECs was measured using a membrane fusion assay adapted from [20]. Briefly, EVs (20 µg/mL protein dose) were labeled with 1 µL of a 1 mM ethanolic solution of Octadecyl Rhodamine B Chloride (R18) for 30 minutes in dark at room temperature in 1 mL of MES buffer (10 mM MES (pH 7.4), 145 mM NaCl, 5 mM KCl)). The unincorporated free R18 dye was removed using NAP-10 columns equilibrated with 1*x* PBS. Next, the EVs were concentrated at 4°C using 3 kD MWCO Amicon Ultra-15 centrifugal filters to remove excess buffer. Ten µg of labeled EVs were added to MES buffer in a thermostated spectrofluorometer FluoroMax-4 (Horiba scientific) and fluorescence was measured continuously at 560-nm excitation and 590-nm emission wavelengths (slit 1.5 nm). Following an equilibration time of 10 min, unlabeled BECs (1 x 10^6^) were added to the EVs, and fluorescence was measured for a further 30 min. The fusion reaction was stopped by the addition of Triton X-100 + octylglucoside at final concentrations of 0.3% and 60 mM respectively, leading to maximal R18 probe dilution. The % fusion data is expressed in terms of % of maximum fluorescence dequenching (**%FD**) = ((F - F_i_)/(F_max_ - F_i_)) x 100, where F is the fluorescence intensity post 30 min of incubation EVs and BECs, F_i_ is the initial fluorescence value of labeled EVs, and F_max_ is the fluorescence intensity followed by detergent-induced membrane disruption.

### 2.13 Statistical analysis

Statistical differences among the mean of control groups and treatment groups or within the treatment groups were measured using one-way or two-way analysis of variance (ANOVA) at 95% confidence intervals using GraphPad Prism 9 (GraphPad Software, LLC). The different levels of significance are indicated as follows: *p<0.05, **p<0.01, ***p<0.001 ****p<0.0001.

## 3. Results and discussion

### 3.1 Physicochemical characterization of BEC EVs and RAW macrophage EVs

#### Throughout the paper, sEVs and m/lEVs are collectively referred to as EVs although we examined them individually

We characterized the particle diameter, dispersity index, zeta potential of BEC EVs and macrophage EVs using dynamic light scattering and determined EV particle concentrations using nanoparticle tracking analysis. BEC-derived sEVs and m/lEVs showed particle diameters of 142.6 ± 2.9 nm and 178.7 ± 3.2 nm, respectively, and showed dispersity indices of 0.38 and 0.27, respectively. Macrophage-derived sEVs and m/lEVs showed particle diameters of 113 ± 0.8 nm and 144.5 ± 7.6 nm, respectively, while both EV subpopulations showed a dispersity index of 0.25 (**Table 1**). These values were consistent with published literature that report EVs as heterogenous particles with sEVs showing particle diameters ranging from 50-200 nm and m/lEVs ranging from 100-1000 nm [6]. BEC-derived sEVs and m/lEVs showed negative zeta potentials of about – 24.2 and −19.5 mV, respectively, and macrophage-derived sEVs and m/lEVs showed negative zeta potentials of about – 19.1and −23.9 mV, respectively. The particle concentration of freshly isolated BEC-derived sEVs and m/lEVs were found to be 9.5 and 7.7 x 10^9^ particles/mL, respectively, and those of macrophage-derived sEVs and m/lEVs were at 2.6 and 1.8 × 10^9^ particles/mL, respectively. The intensity-weighted particle size distribution plots of BEC- and macrophage-derived EVs are presented in **Supplementary Figure 1**.

**Table 1.** Average particle diameters, dispersity indices, and zeta potentials of BEC and RAW macrophage-derived EVs were determined using dynamic light scattering on a Malvern Zetasizer Pro. sEVs and m/lEVs were diluted to a final concentration of 0.1 mg/mL in 1*x* PBS (pH 7.4) for particle size measurements and DI water (pH 7.4) for zeta potential measurements. To measure EV particle concentrations stock samples of sEVs and m/lEVs were diluted in 1*x* PBS (pH 7.4) and analyzed on a multiple-laser ZetaView f-NTA Nanoparticle Tracking Analyzer (Particle Metrix Inc., Mebane, NC). Five 60 s videos were acquired at 520 nm laser wavelengths for particle diameter and concentration measurements. All values were reported as mean ± standard deviation.

|  |  | BEC sEVs | BEC m/IEVs | RAW sEVs | RAW m/IEVs |
| --- | --- | --- | --- | --- | --- |
| **Particle diameter**<br>(nm) | | $142.6 \pm 2.9$ | $178.7 \pm 3.2$ | $113.0 \pm 0.8$ | $144.5 \pm 7.6$ |
| **Dispersity index** | | $0.38 \pm 0.03$ | $0.27 \pm 0.01$ | $0.25 \pm 0.01$ | $0.25 \pm 0.02$ |
| **Zeta Potential (mV)** | | $-24.2 \pm 1.3$ | $-19.5 \pm 0.7$ | $-19.1 \pm 5.2$ | $-23.9 \pm 1.3$ |
| **EV concentration**<br>(particles/mL) | | $(9.5 \pm 1.6) \times 10^9$ | $(7.7 \pm 2.2) \times 10^9$ | $(2.6 \pm 0.5) \times 10^9$ | $(1.8 \pm 1.0) \times 10^9$ |

We identified the presence of CD63 tetraspanin protein in BEC cell lysate and sEVs but not in m/lEVs using western blotting (**Figure 1**). It is important to note that we have not directly compared CD63 expression levels between lysates *vs*. EVs. We note a rather weak expression of CD63 in these sEVs but our future works will employ size exclusion chromatography in combination with sequential centrifugation to further enrich EV markers in the isolated vesicles [29]. We identified the presence CD9 in a pilot experiment in both sEV and m/lEV subpopulations [11, 30–32] using ImageStream flow cytometry (**Figure 2)**. The tetraspanin proteins play a crucial role in the biogenesis or cargo sorting of EVs, explaining their presence in EVs [4]. Nano-sized EVs limit single vesicle analysis and subset specific marker detection using conventional flow cytometry. Scanning and transmission electron microscopy techniques enable the visualization of EV morphology but do not provide a robust quantitative analysis of EM markers. Imaging flow cytometry has demonstrated significant advantages by allowing single vesicle multiparametric analysis, characterization, visualization and functional assessments [33, 34]. Headland *et al.* have demonstrated the use of imaging flow cytometer to analyze microparticle phenotypes. The authors detected the expression of CD66b and CD11b in the neutrophils and their offspring microparticles and highlighted the utility of imaging flow cytometry for extracellular microparticle analysis [35].

**Figure 1.**
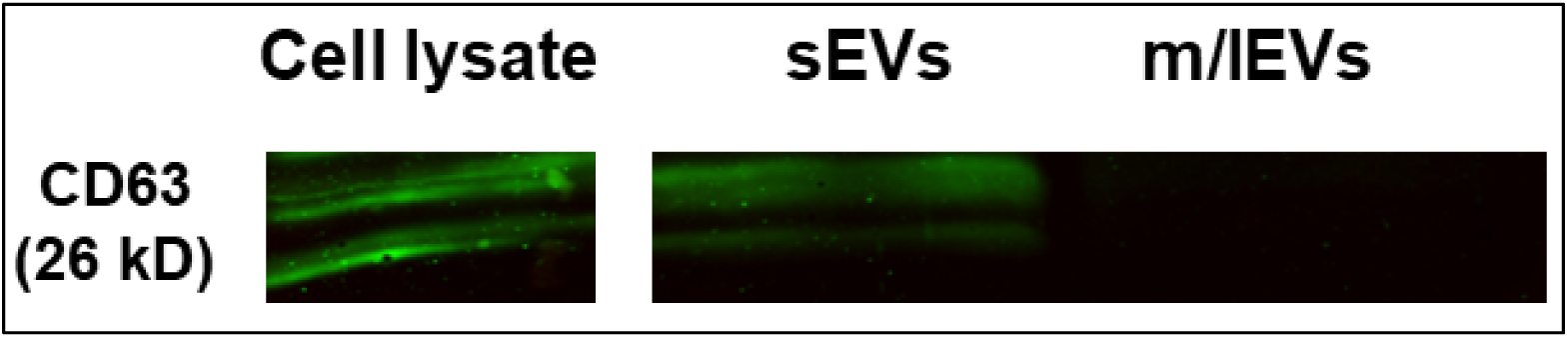
Detection of CD63 in BEC sEVs using western blot analysis. 70 µg protein of each EVs were electrophoresed on a 4–10% gradient sodium dodecyl sulfate-polyacrylamide gel, proteins were transferred onto a nitrocellulose membrane and blocked using blocking buffer. The blot was incubated with CD63 primary antibody overnight at 4°C followed by secondary antibody incubation for one hour at room temperature. The membrane was imaged on 700 nm and 800 nm channels using an Odyssey M imager (LI-COR Inc. Lincoln, NE).

**Figure 2.**
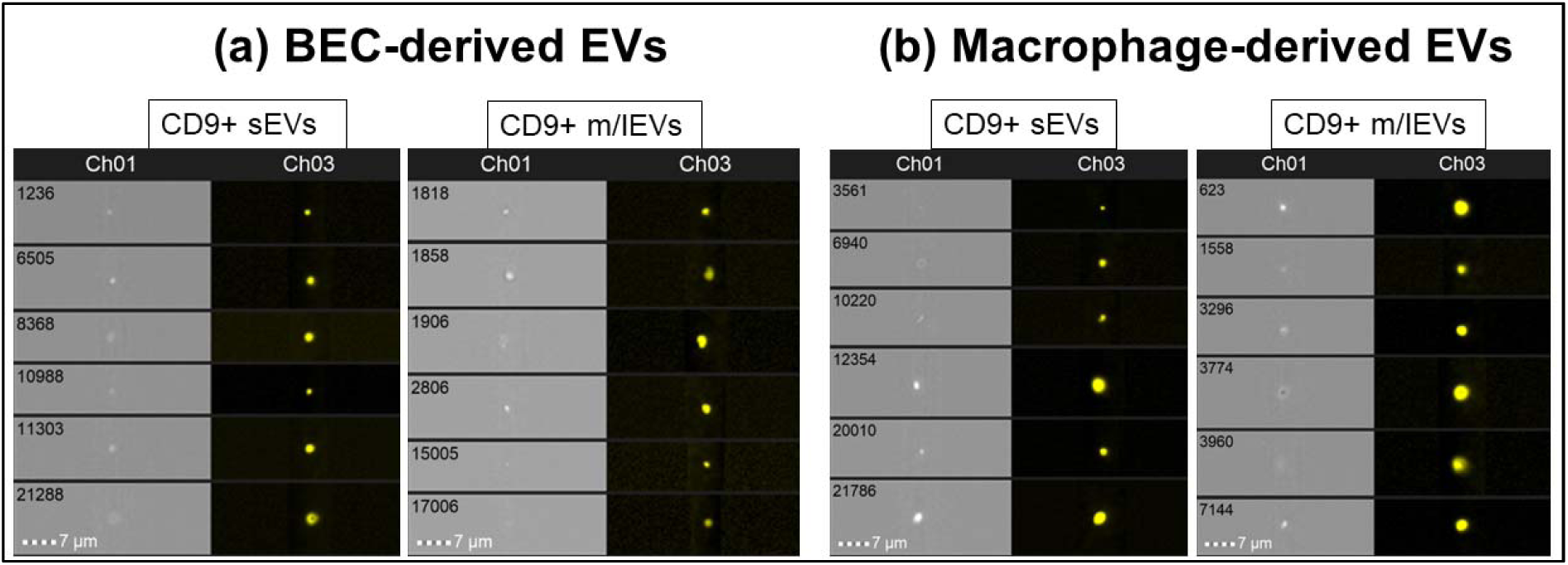
Representative images of CD9-positive EVs. EVs (1x 10^8^ particles/mL) were labeled with anti-CD9 (1 µg) antibody. EVs were incubated with the antibody for 60 minutes at room temperature in the dark. The acquisition of EVs using ImageStreamX was performed on a 488 nm laser at 60× magnification and 7 µm core size. Channels 01 and Ch09 were set to brightfield and channel 03 was used to detect the antibody fluorescence signals.

The goals of this study were two-fold: *first*, we wanted to understand the uptake mechanism involved in the internalization of BEC-derived EVs into the recipient BECs and *secondly*, we wanted to understand the potential effects of homotypic/heterotypic interactions between the EV donor cell-recipient cell type on the uptake of EVs into the recipient cells: specifically, do homotypic interactions between recipient BECs treated with BEC-EVs result in greater levels of cellular uptake compared to recipient BECs treated with macrophage-EVs? We chose a human BEC cell line and RAW 264.7 macrophages and their secreted EVs as our experimental models. BECs are known to have limited pinocytic capabilities [36–38] as opposed to phagocytic cells like RAW 264.7 macrophages. We expected that their discrete differences in endocytic capabilities will allow us to understand any potential differences in the internalization of EVs into the respective recipient cells.

### 3.2 Effect of the labeling dye chemistry on EV uptake into recipient cells

We first determined the effect of the fluorescent dye chemistry used for EV labeling on the uptake of EVs into the recipient cells. We compared the cellular uptake of EVs labeled either with a lipophilic dye, PKH67 or calcein AM ester dye that is cleaved by intravesicular esterases in EVs with intact membranes [27, 39, 40]. We used flow cytometry as a quantitative tool to understand the uptake of EVs into the recipient cells. We used % EV uptake to denote the cell population that is positive for fluorescent EVs as a measure of frequency of EV uptake.

#### 3.2.1 Uptake of BEC EVs into recipient BECs

In the case of recipient BECs treated with 50 µg BEC-EVs (**Figure 3**), we observed ∼10-fold and ∼17-fold increases in the % EV uptake for sEVs and m/lEVs, respectively, for calcein AM- *vs*. PKH67-EV-treated groups (**Figure 3a and 3b**). In the case of recipient RAW macrophages treated with 50 µg macrophage-EVs (**Figure 4**), we observed comparable levels of uptake but small/∼1.2- and 1.6-fold increases in the % sEV and m/lEV uptake, respectively for calcein AM *vs.* PKH67-EV-treated groups (**Figure 4a and 4b**).

**Figure 3.**
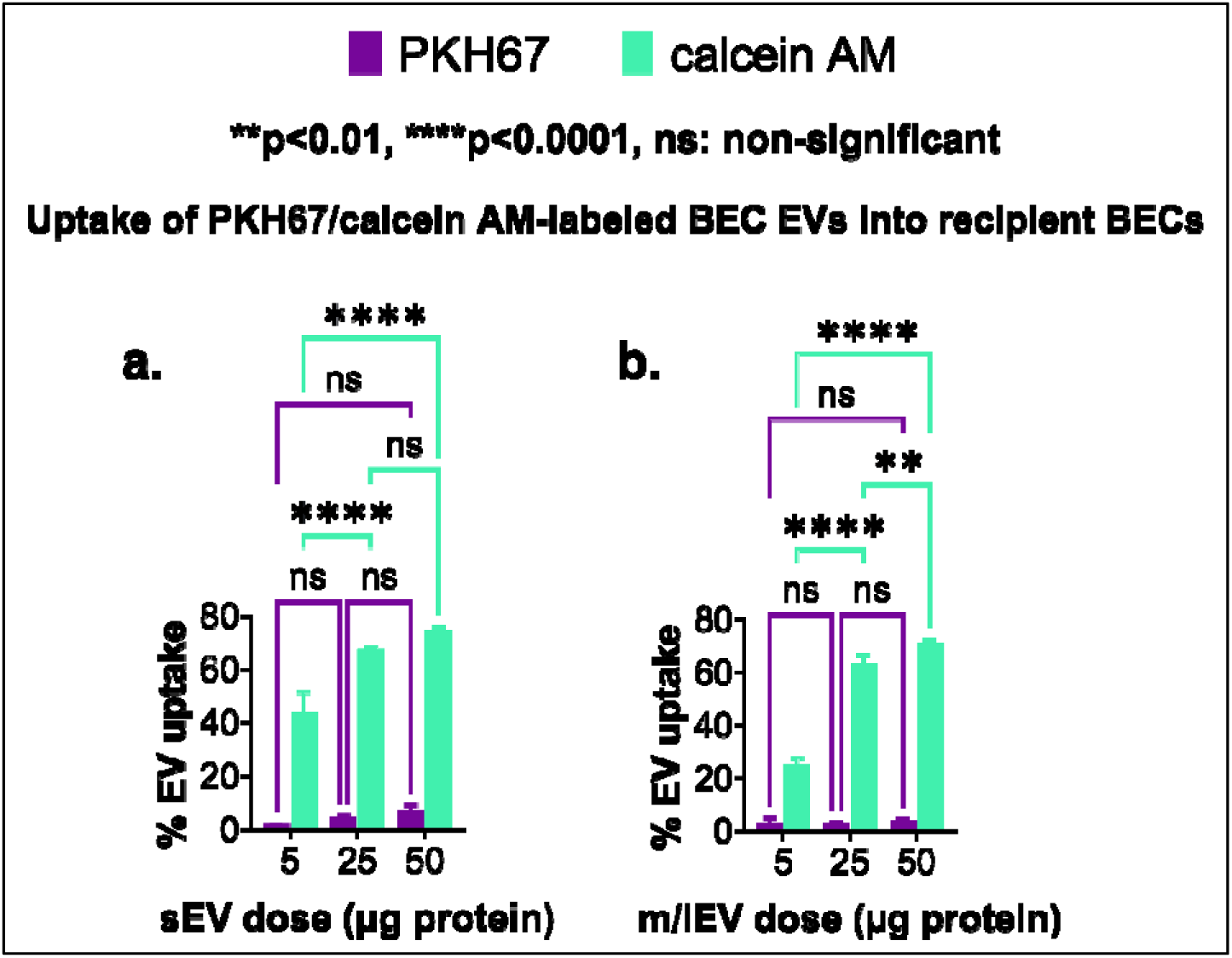
Uptake of BEC-derived EVs into recipient BECs. The recipient BECs were treated with either PKH67- or calcein AM-labeled EVs at 5, 25, and 50 µg EV protein/well for 4 h. The %EV uptake was determined using flow cytometry. Tukey’s multiple comparisons test was used for comparative analysis using two-way ANOVA. Data are presented as mean±SD (n=3).

**Figure 4.**
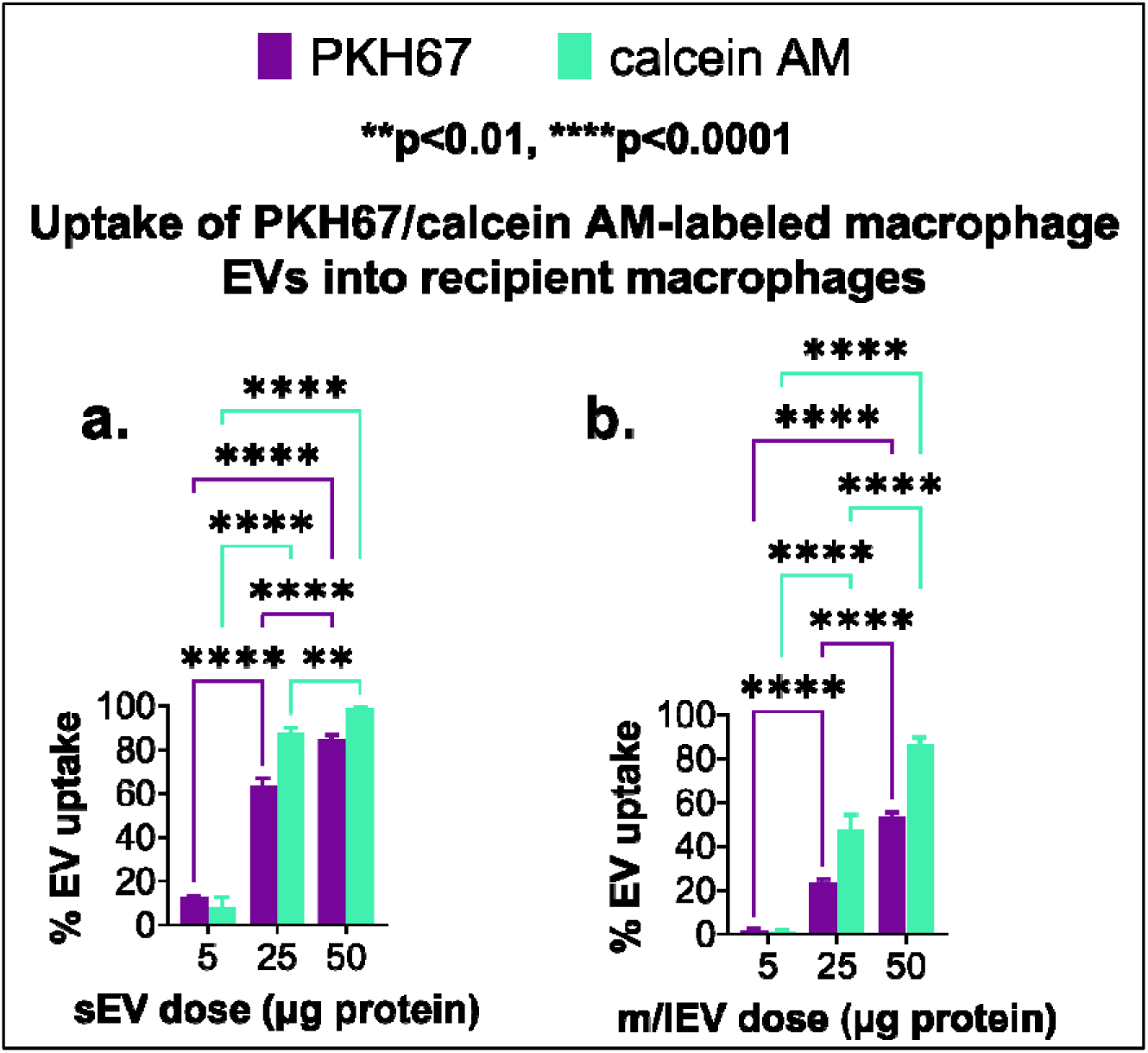
Uptake of macrophage-derived EVs into recipient macrophages. Recipient macrophages were treated with either PKH67- or calcein AM-labeled EVs at 5, 25, and 50 µg EV protein/well for 4 h. The % EV uptake was determined using flow cytometry. Tukey’s multiple comparisons test was used for comparative analysis using two-way ANOVA. Data are presented as mean±SD (n=3).

Our data from **figures 3 and 4** demonstrated two interesting findings. *First*, we noted a dose-dependent increase in the EV uptake only when the recipient BECs were treated with calcein AM-labeled EVs while PKH67-labeled EVs showed dramatically lower levels of %uptake (**Figure 3**), as much as 10-fold lower compared to calcein-labeled EVs at the highest tested dose of 50 µg EV protein. *Second*, in sharp contrast to the data observed with recipient BECs, both PKH67- as well as calcein AM-labeled EVs showed comparable levels of %uptake in the recipient macrophages (**Figure 4**). These observations underscore the importance of choosing appropriate dye labeling chemistry when evaluating EV cellular uptake in different cellular models. The lipophilic PKH67 dye is widely used in EV cellular uptake studies [41]. However, it is known that labeling EVs with a lipophilic PKH dyes can affect the membrane-membrane fusion between EV and recipient cells and also affect the particle diameter of the labeled EVs [42]. It is also known that lipophilic PKH increase the EV particle diameter [43] and concurrently, will affect the cellular uptake of labeled/larger EVs. It is likely that the larger PKH67-labeled EVs are still internalized by the phagocytic macrophages while the low pinocytic BECs do not endocytose such large particles. Calcein AM is a membrane permeable non-fluorescent dye that undergoes hydrolysis by the cytosolic esterases present in the membrane-intact EVs. We and others have demonstrated the suitability of labeling EVs with calcein AM [15, 44, 45] to determine EV membrane integrity under various storage temperatures as well as EV stability post freeze-thaw cycles. Overall, our findings from **figures 3 and 4** allowed us to: *first,* select an optimal EV dose for future experiments; *second,* confirm the suitability of calcein AM for labeling the EVs for tracking the cellular uptake of EVs in low pinocytic BECs as well as macrophages.

In addition to quantifying uptake using flow cytometry, we used epifluorescence microscopy to visualize m/lEV uptake into the recipient cells **(Supplementary Figure 2)**. We observed a dose dependent increase of m/lEV uptake in the calcein AM-labeled EVs as opposed to PKH67 labeled-EV samples. Next, using z-stack images, we visualized the uptake of calcein AM-labeled BEC-derived EVs into recipient BECs. We confirmed the intracellular uptake of EVs using the z-stack fluorescence microscopy and ruled out the possibility that EVs merely stick to the cell surface. Representative images of z-stack in slice view are presented in **Supplementary Figure 3**. Representative flow cytometry histogram plots for figures 2 and 3 are presented in Supplementary Figures 4 and 5.

Following the selection of optimal EV dose and the labeling dye (calcein AM) to measure EV uptake, we next studied the mechanism of EV internalization into recipient cells using endocytosis inhibitors and the effect of homotypic/heterotypic interactions on EV uptake into recipient cells.

### 3.3 Effect of homotypic *vs.* heterotypic interactions on the cellular uptake of EVs in the presence of endocytosis inhibitors

In the experiments below, sucrose, genistein and 5-(N-ethyl-N-isopropyl) amiloride (**EIPA**) were used to block clathrin-mediated, caveolae-mediated and macropinocytic uptake pathways, respectively. The goal of this study was to determine the endocytosis pathways involved in EVs internalization into the recipient cells. It is important to note that endocytosis inhibitors were present throughout the EV exposure period. We used % EV uptake as an output to represent the cell population that is positive for fluorescent signal as a measure of frequency of EV uptake and we used MFI as a measure of extent of EV uptake. **Scheme 1** is a graphical representation of the experimental groups used to study homotypic-heterotypic effects on EV cellular uptake.

**Scheme 1.**
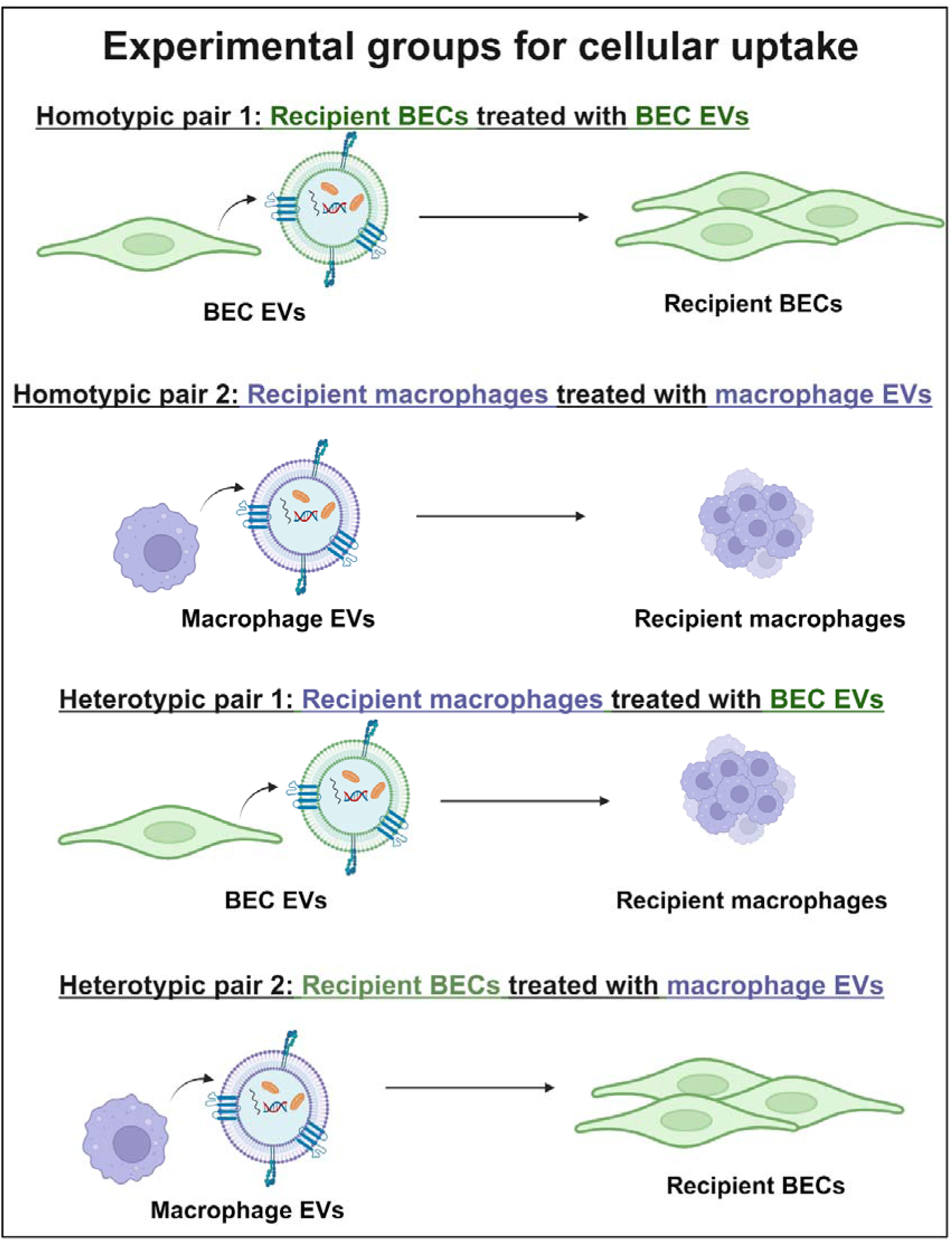
A graphical representation of the homotypic heterotypic experimental groups use d in the cellular up take studies in **Figure 5**.

Comparison of overall uptake/MFI values between recipient BECs and macrophages (naïve uptake/no inhibitors involved) indicated that macrophages show uptake levels that are at least 5-fold greater than those of recipient BECs suggesting that phagocytic macrophages internalize EVs more avidly compared to low pinocytic cells like BECs.

Figure 5a. Homotypic pair 1: Recipient BECs treated with BEC EVs

**Figure 5.**
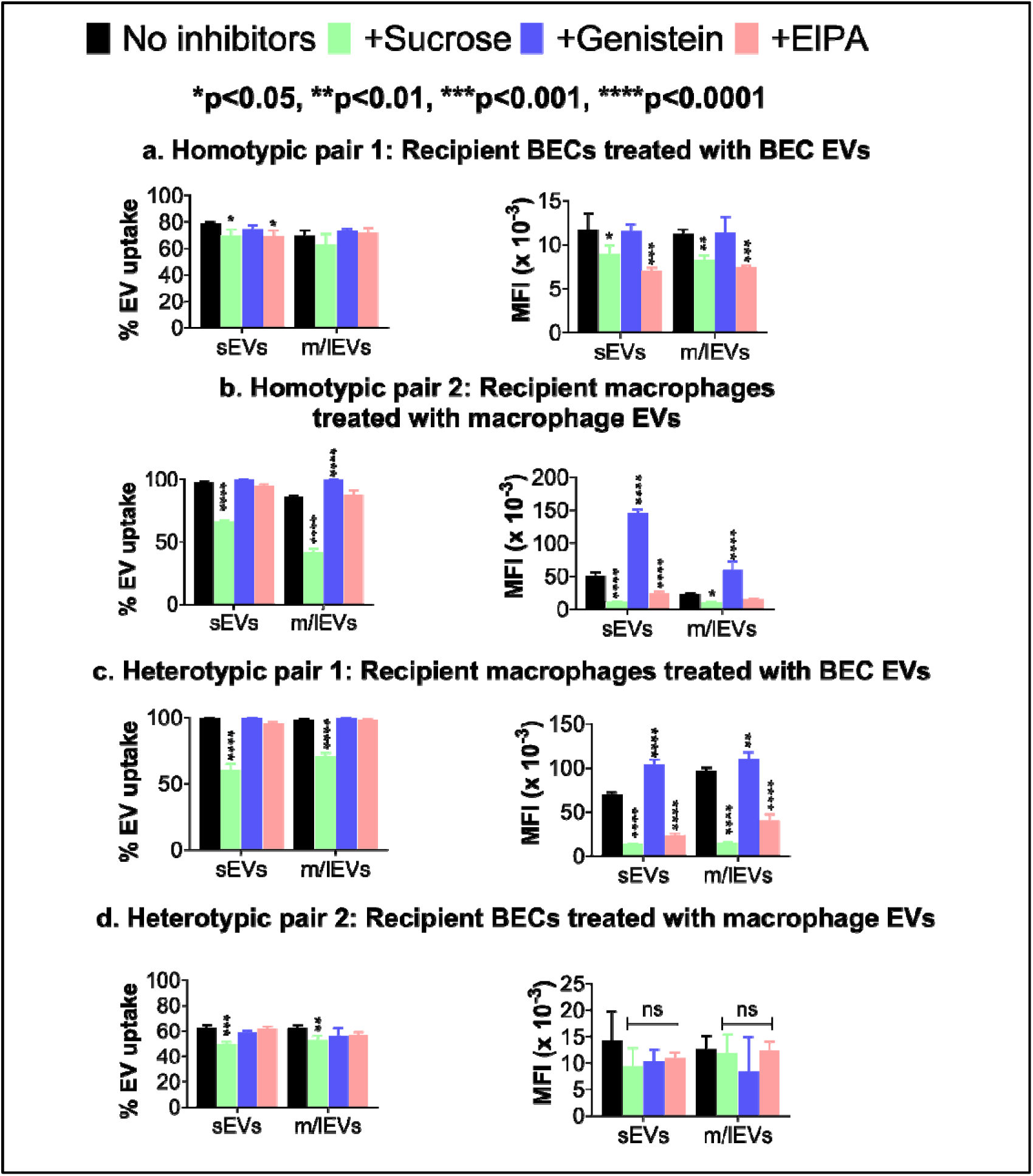
Effect of endocytic inhibition on the cellular uptake of EVs. Recipient cells were pre-incubated for 0.5 h with endocytosis inhibitors and then were treated for 4 h m/lEVs and sEVs at 25 µg (recipient BECs) and 50 µg protein (recipient macrophages)/well in the presence of fresh endocytosis inhibitors. The % EV uptake and MFI values were determined using flow cytometry. Dunnett’s multiple comparisons test was used for statistical analysis in comparison with the control/‘no inhibitors’ group. Data are presented as mean±SD (n=3).

Recipient BECs showed %uptake levels varying from 70 to 80% when treated with BEC-EVs. When clathrin and macropinocytosis pathways were blocked, sEV-treated cells showed a significant reduction in %uptake and MFI values whereas, m/lEV-treated cells showed only a significant reduction in MFI values while the %uptake remained unaffected.

<u>Figure 5b. Homotypic pair 2: Recipient macrophages treated with macrophage EVs</u>

Recipient macrophages showed %uptake levels varying from >80 to 100% when treated with macrophage-EVs. Both sEV- and m/lEV-treated cells showed a significant reduction in %uptake and MFI values when clathrin-mediated endocytosis was blocked using sucrose. sEV-treated cells also showed a significant reduction in the MFI values but no changes in %uptake was noted when macropinocytosis was inhibited using EIPA. Interestingly, we observed an increase in the uptake of both subtypes of EVs when caveolae-mediated endocytosis was blocked using genistein. Overall, we observed that recipient macrophages showed reduced EV uptake when endocytic uptake was inhibited while the less pinocytic BECs remained mostly unaffected.

<u>Figure 5c. Heterotypic pair 1: Recipient macrophages treated with BEC EVs</u>

Recipient macrophages showed nearly 100% uptake levels when treated with BEC-EVs. Both sEV- and m/lEV-treated cells showed a significant reduction in %uptake and MFI values when clathrin-mediated endocytosis was blocked using sucrose. Both sEV- and m/lEV-treated cells also showed a significant reduction in MFI values when macropinocytosis was blocked using EIPA. Similar to homotypic pair 2 (macrophage EVs➔recipient macrophages), we observed an increase in the uptake of both subtypes of EVs when caveolae-mediated endocytosis was blocked using genistein. As noted earlier with homotypic pair 2, we observed the higher sensitivity of recipient macrophages to endocytic inhibition in comparison to the less pinocytic BECs.

<u>Figure 5d. Heterotypic pair 2: Recipient BECs treated with macrophage EVs</u>

Recipient BECs showed nearly 60% uptake levels when treated with macrophage-EVs. Both sEV- and m/lEV-treated cells showed a significant reduction in %uptake when clathrin-mediated endocytosis was blocked using sucrose. However, we did not note a reduction in MFI values.

Our data from Figure 5 suggest that although there were no significant reduction in the %uptake values, the MFI values were significantly lower in different experimental groups indicating that although the *percent of cells internalizing EVs remain the same, the extent/intensity of EV uptake decreased significantly as a function of endocytic inhibition* (the caveolae pathway exception in **Figures 5b and 5c** is discussed below). Overall, our data (Figure 5) shows a greater level of uptake (∼70-80%) when recipient BECs were treated with BEC-derived EVs (Figure 5a) in contrast to recipient BECs treated with macrophage-derived EVs (∼60%) (Figure 5d). These observations suggest that homotypic interactions allow improved uptake into recipient BECs compared to heterotypic interactions, confirming our hypothesis. Recipient macrophages internalized both BEC-derived and macrophage-derived EVs to comparable extents, suggestive of their phagocytic potential—irrespective of homotypic *vs*. heterotypic interactions. Representative histogram plots for Figure 4 are presented in Supplementary Figure 6.

Interestingly and unexpectedly, recipient macrophages treated with macrophage EVs or BEC EVs showed *dramatically increased the extent of EV uptake/MFI* (**Figure 5b and 5c)** when caveolae-mediated endocytosis was inhibited using genistein. These increases were observed only with the recipient macrophages but the recipient BECs remained unaffected by caveolae inhibition. Genistein is a tyrosine-kinase inhibitor that disrupts the local actin network at membrane sites and inhibits the recruitment of dynamin II, two known critical events in caveolae-mediated uptake mechanism. Dynamin II is required for scission of the caveolae membrane pit to caveolae vesicle [46, 47]. We believe that a small fraction of EVs might be deposited in the formed caveolae pits but still are unable to be internalized because genistein inhibits the dynamin recruitment, preventing scission of caveolae pit into the vesicle. We speculate that this phenomenon might be manifesting as an increased fluorescence intensity or “apparent increase in the extent of EV uptake” as compared to the untreated cells. In short, while genistein-mediated blockage prevented complete internalization of EVs, the added EVs still likely bound to the recipient cells and stayed inside of the caveolae pits manifesting as EV fluorescent signals **(Scheme 2)**. Alternatively, we speculate that cells may have upregulated other endocytic pathways to compensate for the inhibition of caveolae mediated uptake—and ultimately increase EV internalization. It is also likely that blocking of caveolae mediated pathway using genistein treatment may have increased the membrane fluidity, enhancing the EV uptake through other pathways [17]. Tian *et al.* observed similar findings when they pre-treated rat pheochromocytoma PC12 cells with 5–50 µM nystatin (alternate pharmacological inhibitor of caveolae-mediated endocytosis). Following nystatin exposure, recipient PC12 cells were incubated with PC12 cell-derived exosomes and uptake of EVs was studied using confocal microscopy and quantified using relative fluorescence intensities. The authors noted ∼3 fold increase in EV uptake in 50 µM nystatin-treated cells as compared to control/no inhibitor group. In contrast to recipient macrophages, the lack of sensitivity of recipient BECs to genistein-mediated increases in apparent uptake/MFI may be explained by the overall lower pinocytic potential of BECs in comparison to professional phagocytes like macrophages.

**Scheme 2.**
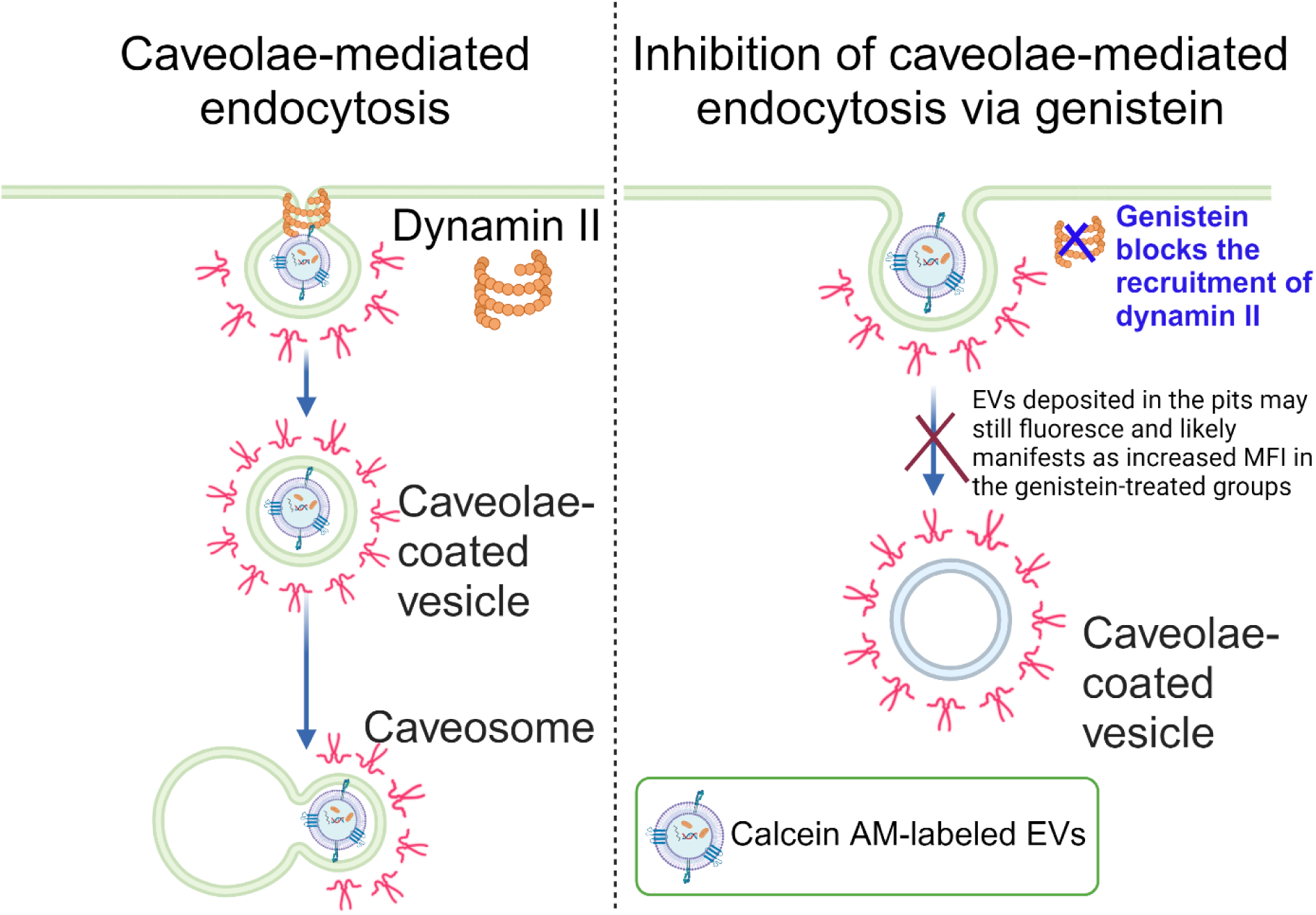
Speculative hypothesis explaining the “apparent” increase in MFI in genistein-treated EV groups in Figures 4b and 4c *despite* the known effects of genistein-mediated inhibition of caveolae endocytosis.

The size of the nanoparticles play a significant role in the route of internalization. The caveolae-coated vesicle can accommodate particles in the range of ∼50-100 nm, thereby limiting the entry of the major fraction of EVs above this size [24, 48–50]. As shown in **Table 1**, both BEC and RAW macrophage-derived EV subpopulations have average particle diameters >100 nm. On the other hand, clathrin-dependent endocytosis internalizes particles with a reported upper size limit of ∼200 nm [51, 52] and macropinocytosis can internalize materials even >1 µm in size [47].

Our data suggesting the involvement of clathrin-dependent and macropinocytosis pathways in the internalization of EVs but the lack of caveolae dependence is in contrast with results reported by other groups. For example, clathrin-dependent endocytosis was not a reported uptake pathway when recipient HeLa (human cervical carcinoma) cells were treated with A431 (human epidermoid carcinoma) cell-derived EVs [24]. Also, we observed a significant reduction in EV uptake when recipient BECs treated with macrophage-derived EVs *only* in the presence of sucrose (inhibitor of clathrin-dependent endocytosis) unlike Yuan *et al*. [16], where the authors had demonstrated the reduction in macrophage-derived exosome uptake into recipient BECs in the presence of inhibitors of *all* three endocytosis pathways (clathrin-dependent, caveolae-mediated and macropinocytosis). Our data, however, agrees with reports that demonstrated the involvement of clathrin-mediated and macropinocytosis but hinted at the lack of involvement of caveolae-mediated endocytosis for EV internalization [22, 25]. Tian *et al.* demonstrated that rat pheochromocytoma PC12 cell-derived EVs were internalized into the PC12 cells via clathrin and macropinocytosis pathways. Ilahibaks *et al*. showed clathrin-dependent endocytosis and macropinocytosis-mediated delivery of human embryonic kidney 293FT (HEK293FT)-derived EVs into T47D stoplight reporter cells [22, 25]. Overall, these discrepancies highlight the importance of *<u>not</u>* generalizing pathways of EV uptake into all types of cells as the EV donor species-recipient interactions seem to play a key role in determining the EV uptake pathway.

### 3.4 Effect of heparin on EV uptake

Given the minimal role of endocytosis pathways in the uptake of EVs into recipient BECs, we sought to determine the involvement of heparan sulfate proteoglycans (**HSPGs**) in the uptake of EVs. We narrowed down our focus to recipient BECs in this study as these low pinocytic cells are the model of our interest in the context of our previously published works on EV mitochondria delivery to the blood-brain barrier [12–15]. We treated the recipient BECs with BEC-derived EVs in presence or absence of heparin for either 30 minutes or four hours. HSPGs are located on the cell surface and the extracellular matrix and are composed of covalently attached HS chains, a type of glycosaminoglycan [53, 54]. HSPGs are abundantly present in the brain endothelium [55–57] and are known to be involved in the receptor-mediated endocytosis of HIV-1 particles [57, 58] and HIV-TAT protein [59]. Many reports have demonstrated that EVs from cancer cells [60–62] interact with HSPGs and the role of the HSPG inhibition on the effects of EV-HSPG interactions. Atai *et al.* demonstrated involvement of HSPG receptors in the attachment and internalization of EVs derived from U87/GBM 11/5 and 293T/HUVEC into the respective recipient cells [21].

Assuming that heparin blockade of HSPG receptors in recipient cells would inhibit dynamin-dependent endocytosis [55] and macropinocytosis [63] pathways, our goal was to determine if HSPG blockade may have an effect on the uptake of EVs into the recipient BECs. The internalization/adherence of EVs have been observed as early as 30-minutes post EV exposure to the recipient cells [64, 65]. Therefore, we chose the 30-minute EV-recipient cell incubation to determine whether heparin blocks the early/acute phase of EV uptake into the recipient cells. Our results show that heparin shows no effect of EV uptake regardless of the exposure time (Figure 6), suggesting that EVs may be internalized into recipient BECs primarily via membrane fusion.

**Figure 6.**
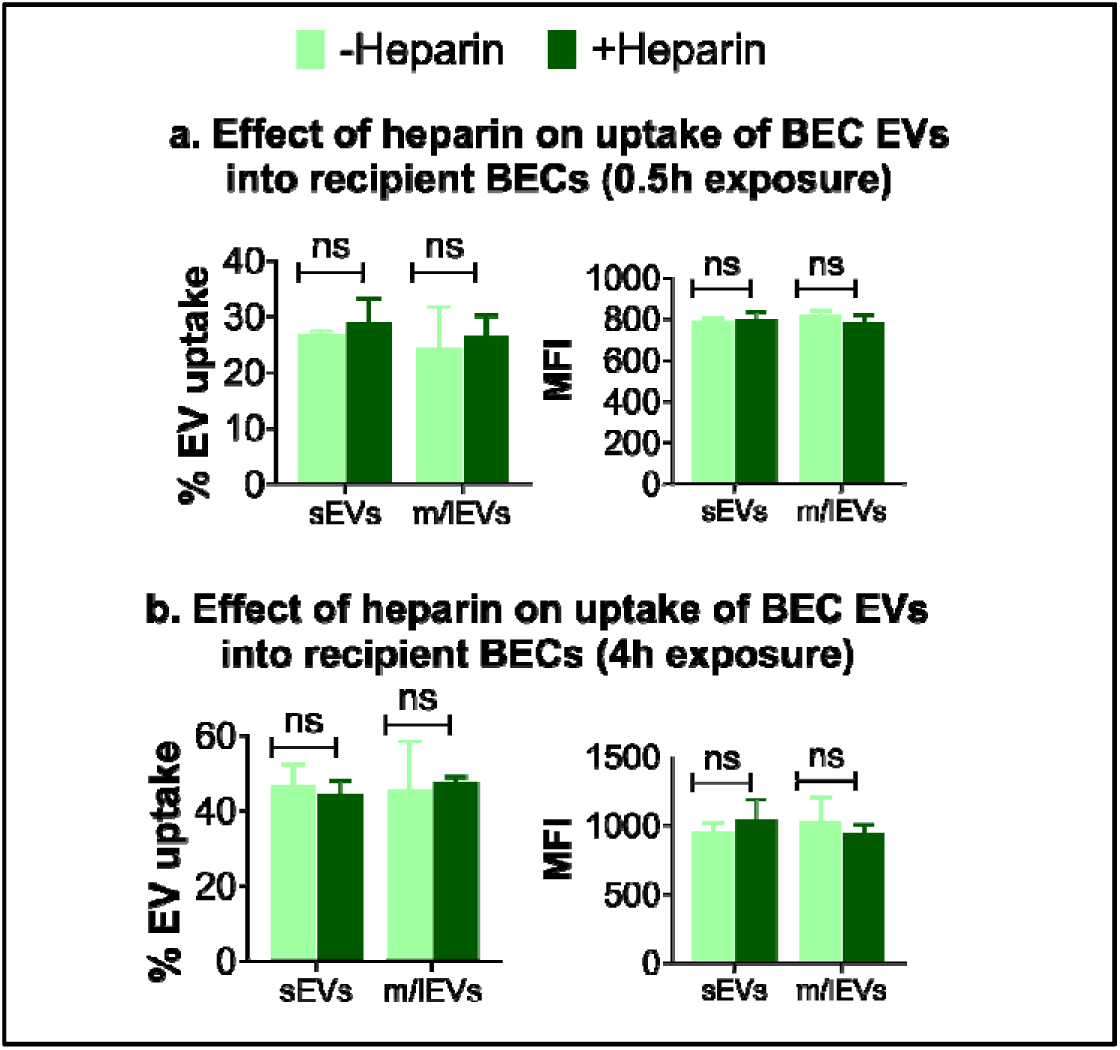
Effect of heparin on EV uptake in recipient BECs. Recipient BECs were treated for 0.5 h with heparin (100 µg/mL) or 1*x* PBS along with BEC derived sEVs and m/lEVs at 25 µg protein/well (**a**). Recipient BECs were pre-incubated for 0.5 h with heparin (100 µg/mL) or 1*x* PBS and then were treated for 4 h with fresh heparin (100 µg/mL) or 1*x* PBS along with BEC-derived sEVs and m/lEVs at 25 µg protein/well (**b**). The % EV uptake and MFI values are determined using flow cytometry. Šídák’s multiple comparisons test was used to determine statistical significance, if any. Data are presented as mean±SD (n=3).

Interestingly, we observed no significant changes in EV uptake irrespective of presence or absence of heparin at 0.5 h *vs.* 4 h (Figure 6). These data seem to align well with our pharmacological inhibition studies (**Figure 5a and 5d**) that showed that blocking of key endocytosis pathways did not seem to significantly affect the extent of EV cellular uptake in the recipient BECs. Interestingly, we observed a dramatic decrease in the uptake of PKH67-labeled EVs into the recipient BECs compared to the greater levels observed in the case of calcein AM-labeled EVs (Figure 3). We speculate that labeling EVs with a lipophilic dye like PKH67 can affect the EV size and therefore its fusion with recipient cell membranes [42, 43] (Figure 3) suggesting the involvement of membrane fusion as a primary route of EV uptake in low pinocytic cells such as BECs. The homotypic interactions between the lipidic EV membranes and the recipient BEC membranes may drive the passive EV transport via membrane fusion. Hence, we studied the fusion of EVs with the recipient BECs using a self-quenching R18 probe-based membrane fusion assay.

We measured the fusion activity of EVs with BECs using a lipid mixing fusion assay adapted from Parolini *et al*. [20]. Hoekstra *et al.* developed a R18 fluorescence probe-based fusion assay to measure the kinetics of fusion between biological membranes [66]. The authors studied the fusion of Sendai virus with erythrocyte ghosts using a fusion assay [66, 67]. R18 probe is a self-quenching dye where the self-quenching phenomenon reduces the fluorescence intensity of the probe when the fluorophore molecules are in close proximity [68]. R18 undergoes self-quenching when incorporated in high surface density lipid membranes (here, EVs). This assay is based on the reversal of fluorescence self-quenching of R18 probe, also termed as “dequenching”. Fusion between the biological membranes (EVs and BECs) causes reduced surface density, leading to dilution of the R18 probe and ultimately resulting in dequenching of R18 probe and increased fluorescence intensity (as a consequence of reversal of self-quenching) [66]. We noted a gradual increase in the fluorescence intensity over a course of 30 minutes upon incubation of R18-EVs with unlabeled BECs (**Figures 7a and 7b**). It is important to note that the control (no BECs added/EVs only) group F_max_ values were nearly two-fold greater for m/lEVs over sEVs—likely indicating that the R18 probe labeling efficiency of EVs varies as a function of lipid composition of EVs. Given that the biogenesis pathways of sEVs *vs.* m/lEVs are different [4, 7, 69, 70], it is fair to expect that their lipid composition will also be different. Our future studies will involve a thorough comparison of m/lEV *vs*. sEV membrane composition using a lipidomics approach [71] that may shed light on the potential differences in their lipid composition and thereby, differences in R18 labeling efficiencies. The data demonstrates lipid mixing activity of EVs and BEC membranes reaching 15-20% dequenching of the maximum fluorescence obtained after addition of Triton X-100 + octylglucoside detergent mixture (Figure 7c). We observed no spontaneous dequenching of R18-EVs until addition of the detergent mixture indicating the specificity of the measured fluorescence dequenching. Parolini *et al.* demonstrated that the recipient melanoma cells internalize melanoma cell-derived exosomes via membrane fusion using a R18 probe-based membrane fusion assay [20]. Over the course of the 30-minute incubation of R18-EVs and recipient melanoma cells, the authors observed a continuous lipid mixing/dequenching reaching ∼30% of the maximum fluorescence. We also performed a membrane fusion assay to assess the fusion activity of macrophage EVs with recipient macrophages and measured fluorescence dequenching values were as low as <4%, suggesting that macrophages do not utilize membrane fusion process to internalize EVs (**Supplementary figure 7**).

**Figure 7.**
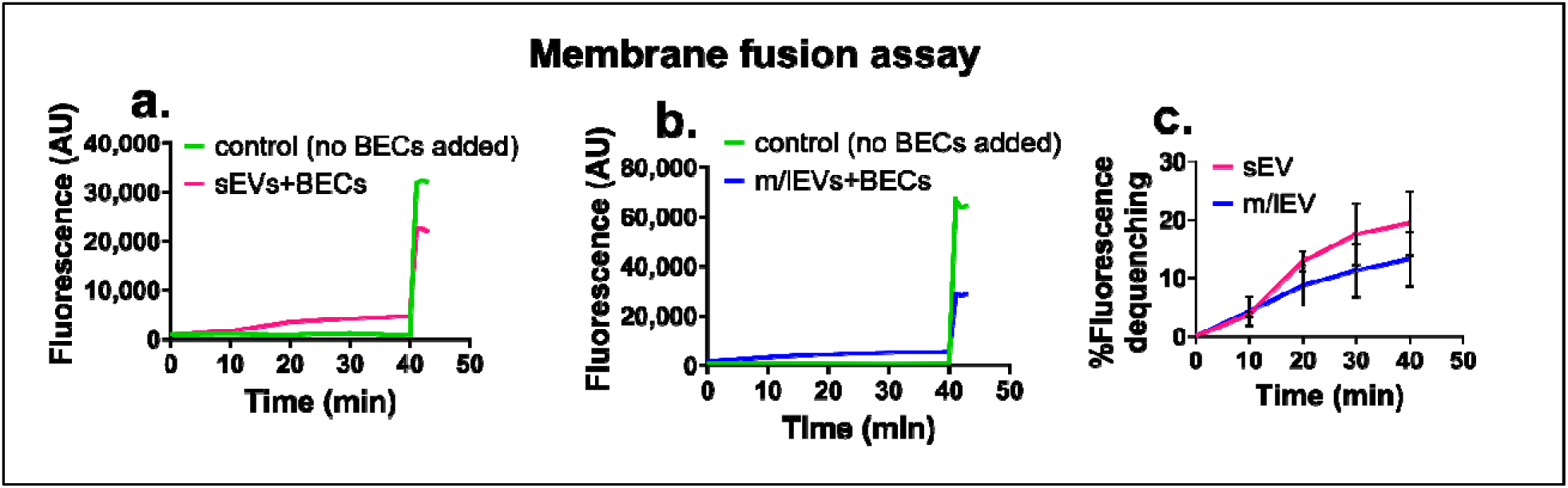
EV-recipient BEC membrane fusion assay. Ten µg of R18-labeled EVs were added to MES buffer in a thermostated spectrofluorometer FluoroMax-4 and fluorescence wa measured continuously at 560-nm excitation and 590-nm emission wavelengths (slit 1.5 nm). Post 10 min of equilibration, unlabeled BECs were added to R18-EVs, and fluorescence wa measured for a further 30 min. The fusion reaction was stopped by the addition of 0.3% Triton X-100 + 60 mM octylglucoside. **(a, b).** R18 labeled sEVs or m/lEVs were left untreated (control) or mixed with unlabeled 1×10^6^ BEC cells (sEV/m/lEV-treated). **(c)**. Fluorescence dequenching (FD) plot where the % FD is defined as = ((F - F_i_)/(F_max_ - F_i_)) x 100. Data is presented as mean±SD (n=3).

In summary, our data demonstrates that the first 30-minutes/early phase of EV uptake into recipient BECs proceeds via membrane fusion and it is likely that endocytic pathways such as clathrin-mediate endocytosis kicks in at later times (Figure 5). The slow kinetics (∼15-20% fluorescence dequenching) of membrane fusion-based uptake likely explains our previous observations showing mitochondria transfer from m/lEVs into recipient BECs takes at least 72 h while limited transfer was noted at the 24- and 48 h time points [15]. Overall, our results highlight the importance of *not* generalizing EV uptake as a process that proceeds via endocytic mechanisms and also underscores the importance of determining uptake routes into low pinocytic cell models such as BECs—given our longstanding interest in the delivery of therapies to protect the BBB [12, 13, 15, 72, 73].

## 4. Conclusions

Overall, our data demonstrated that while blocking clathrin-mediated endocytosis and macropinocytosis pathways slightly reduced the intensity of EV uptake in the recipient BECs, inhibition of these pathways did not completely block EV uptake. Inhibition of the HSPG receptors in recipient BECs demonstrated no effect on the uptake of EVs. This suggests a minor role of endocytosis for EV internalization in the recipient BECs. The membrane fusion assay suggests that the early phase uptake of BEC-derived EVs into recipient BECs proceeds using an energy-independent, membrane fusion processes. Our results provide important insights on the EV uptake mechanisms and also underscore the importance of carefully choosing labeling dye chemistry for studying EV uptake into low pinocytic recipient cells such as BECs.

## Acknowledgements

We are grateful to Dr. Rehana K. Leak and her lab for assistance with using the Olympus microscope.

## Statements and Declarations

### Ethics approval and consent to participate

The study does not involve any animal or human data and so, neither ethics approval nor consent to participate was required.

### Consent for publication

Not applicable (see above)

### Author Contributions

Devika S Manickam conceived the study and its design. Experiments, data collection and analysis were performed by Jhanvi R. Jhaveri, Purva Khare, Paromita Paul Pinky, Jadranka Milosevic, Nevil Abraham and Devika S Manickam. Experimental support and technical insights were provided by Yashika S. Kamte, Manisha N. Chandwani, Paromita Paul Pinky, Si-yang Zheng, and Lauren O’Donnell. The manuscript was written by Jhanvi R. Jhaveri, Purva Khare and Devika S Manickam.

### Data availability

The data generated during the current study will be available from the corresponding author on request.

## List of non-standard abbreviations

BECs: brain endothelial
cells calcein AM: calcein acetoxymethyl ester
EIPA: 5-(N-Ethyl-N-isopropyl) amiloride
EVs: extracellular vesicles
HSPG: heparan sulfate proteoglycans
MFI: median fluorescence intensity
m/lEVs: medium-to-large
EVs sEVs: small EVs

