## Supplementary Material for "Effect of homotypic *vs*. heterotypic interactions on the cellular uptake of extracellular vesicles"

**Particle size distribution of BEC- and macrophage-derived EVs**

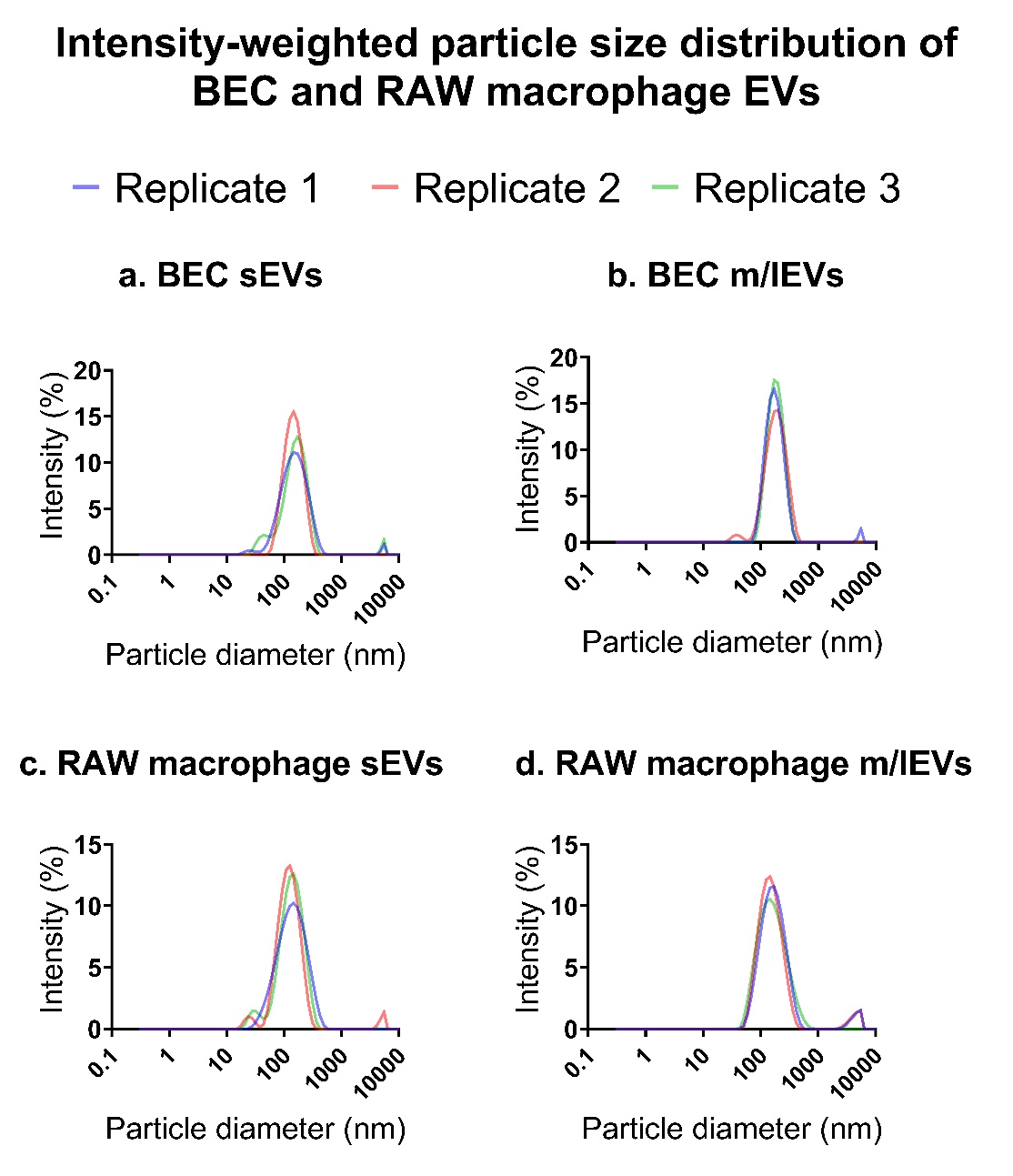

**Supplementary figure 1. Intensity-weighted particle size distribution of BEC and macrophage-derived sEVs and m/lEVs using dynamic light scattering demonstrating the distribution of particle diameters as a function of % scattered light intensity.**

**Uptake of macrophage m/lEVs into recipient macrophages using fluorescence microscopy**

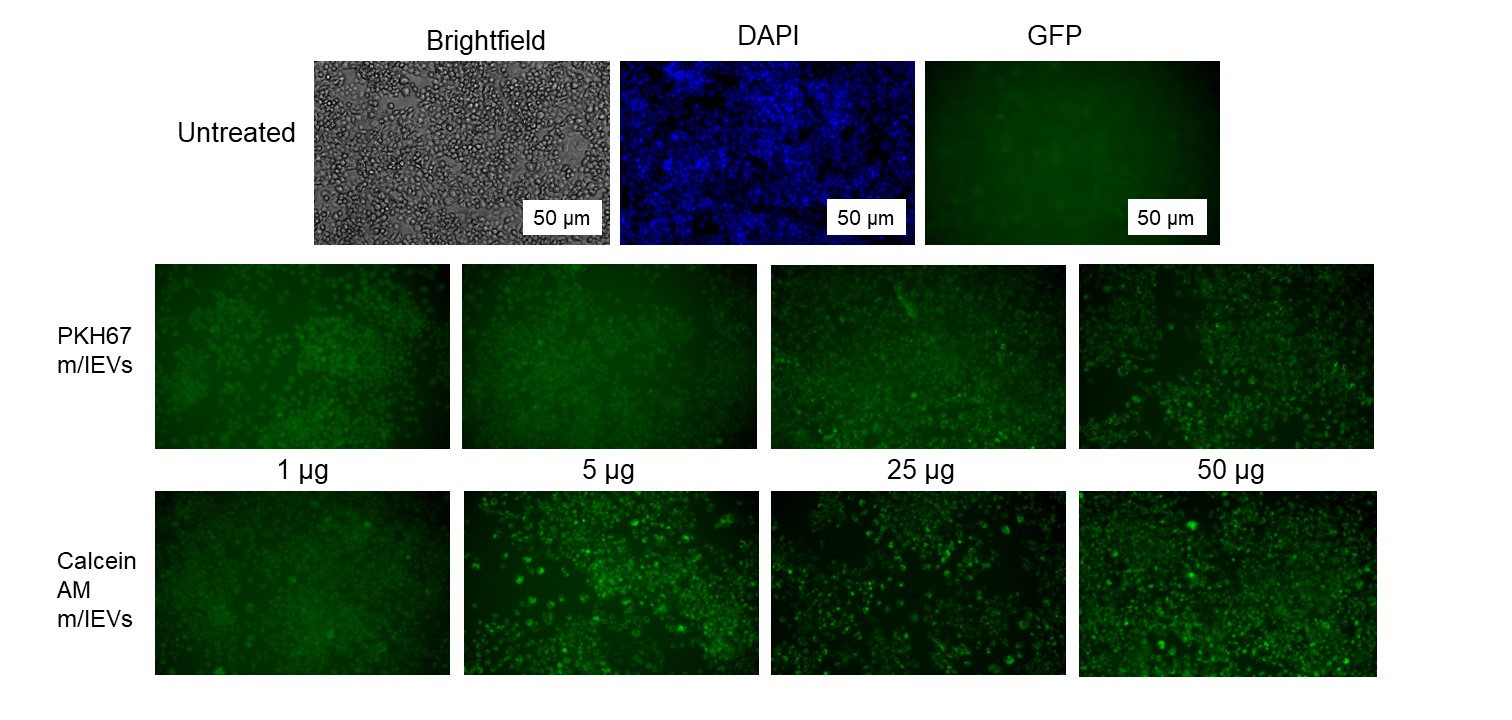

**a.**

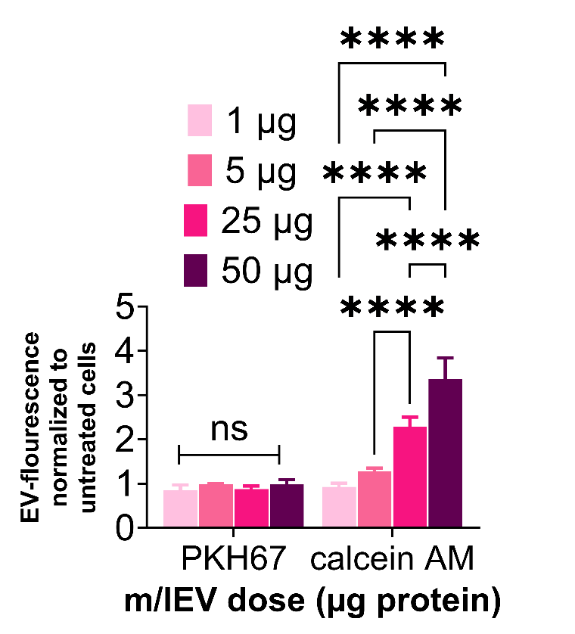

**b.**

**Supplementary Figure 2.** Uptake of macrophage-derived m/lEVs labeled with PKH67 or calcein AM into the recipient macrophages was detected under the GFP channel using an Olympus epifluorescent microscope at 20*x* magnification. Scale bar: 50 µm (**a**). The sum of grayscale signal intensities in the GFP channel were estimated using Olympus CellSens software. The measured intensities were normalized with those of the untreated cells (n=3) (**b**).

Recipient macrophages were incubated with 1, 5, 25 and 50 µg protein dose of calcein AM- or PKH67-labeled m/lEVs. The data suggested a dose dependent increase in calcein AM-labeled m/lEVs while PKH67-labeled EVs showed comparable signals at all doses.

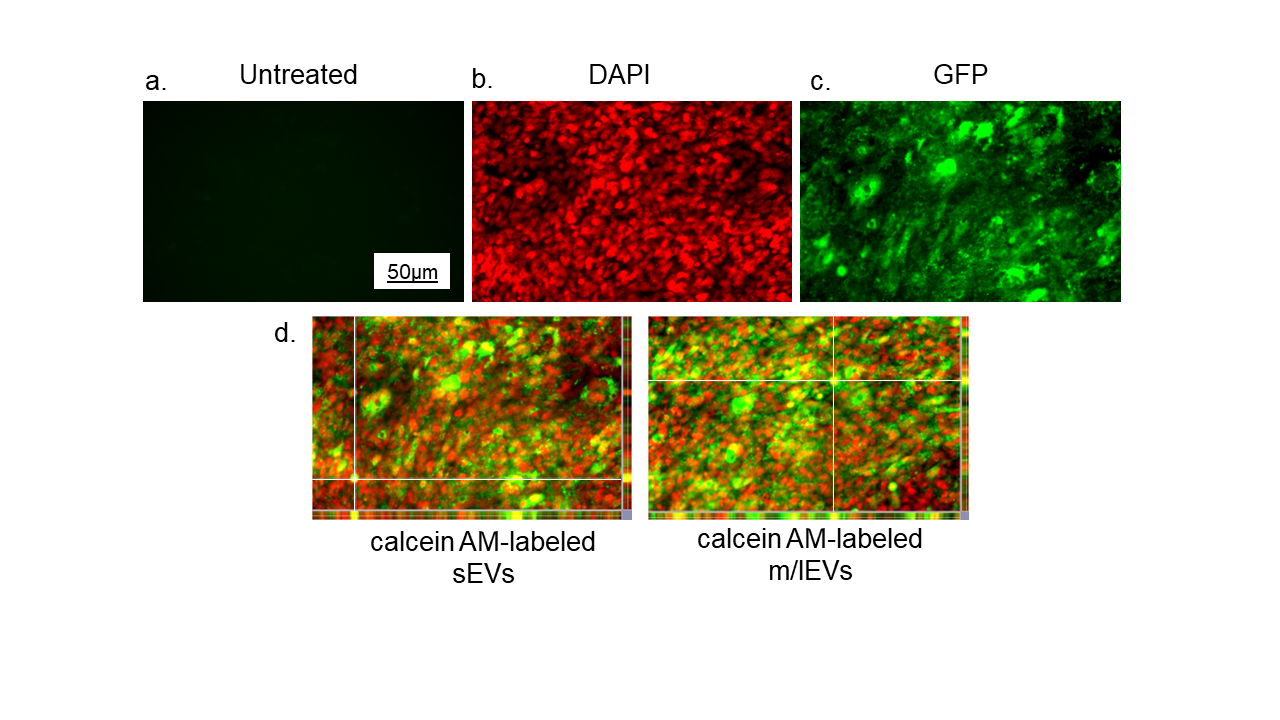

**Supplementary Figure 3. Uptake of calcein AM-labeled EVs (25 µg EV protein) into the recipient BECs using fluorescence microscopy.**

Intracellular EV signals were observed under an Olympus IX 73 epifluorescent inverted microscope using GFP channel (green puncta) at 20x magnification. Intracellular Hoechst labelled nucleus signals were observed using DAPI channel (false colored in red) at 20x magnification (a, b, c). Scale bar: 50 µm. Representative z-stack-slice views were obtained under GFP and DAPI channel using z-stack arrangement (d).

Recipient BECs were treated with 25 µg protein dose of calcein AM-labeled sEVs and m/lEVs for 24 h. Z-stack fluorescence images suggested that EVs were present intracellularly and ruled out the possibility that EVs merely stick to the cell surface. The appearance of “yellow” overlay signals points to the intracellular presence of EVs in the recipient BECs.

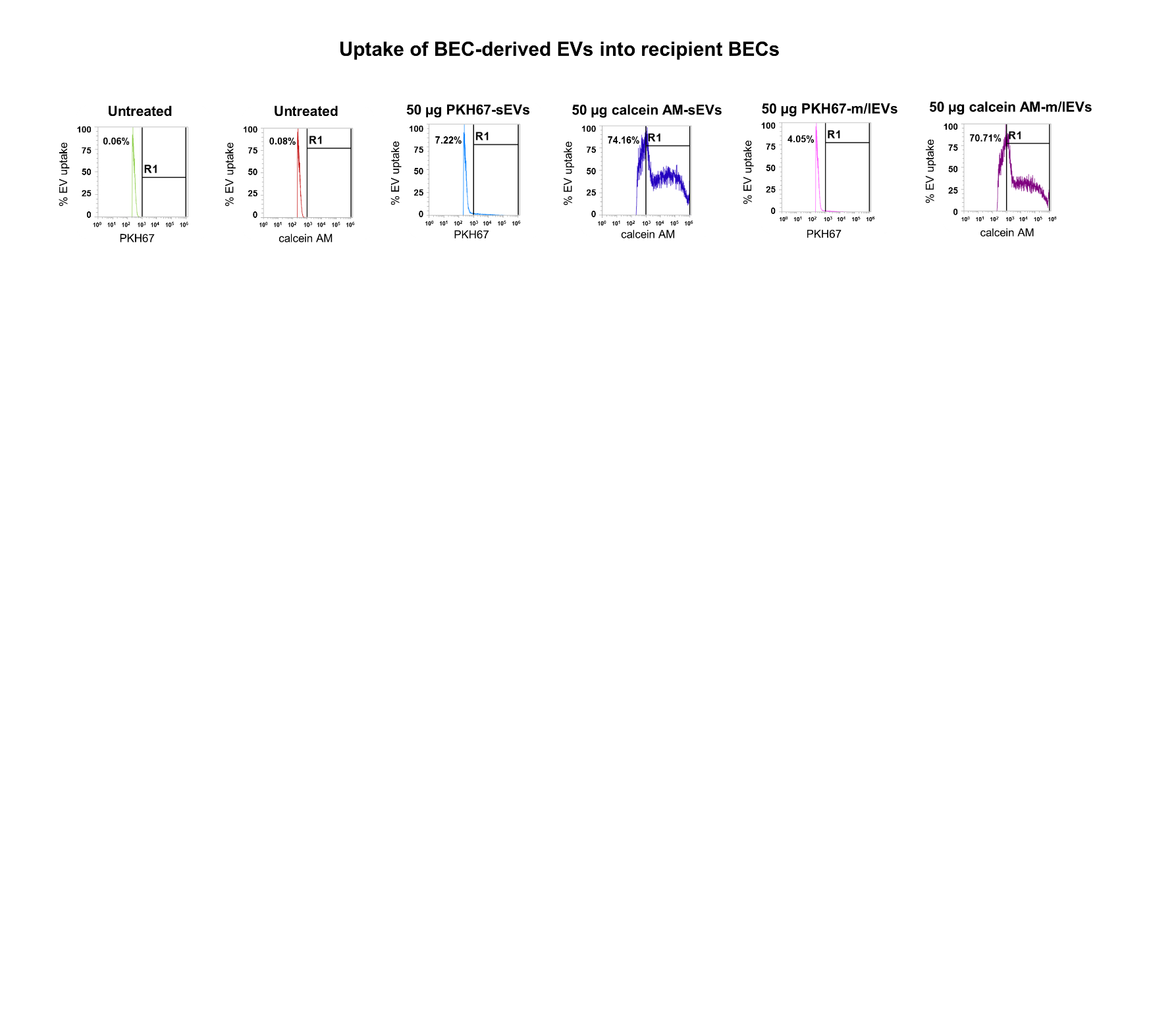

**Supplementary Figure 4. Representative histogram plots showing gating strategy for determining BEC-derived EV uptake into recipient BECs for figure 3 in main text.**

Recipient BECs were treated with either PKH67- or calcein AM-labeled EVs at 5, 25, and 50 µg EV protein/well for 4 h. Data represent PKH67/calcein AM (+) cells noted by the shift past the solid black line after gating the autofluorescence of the untreated cells. The histogram plots are representative of triplicate samples.

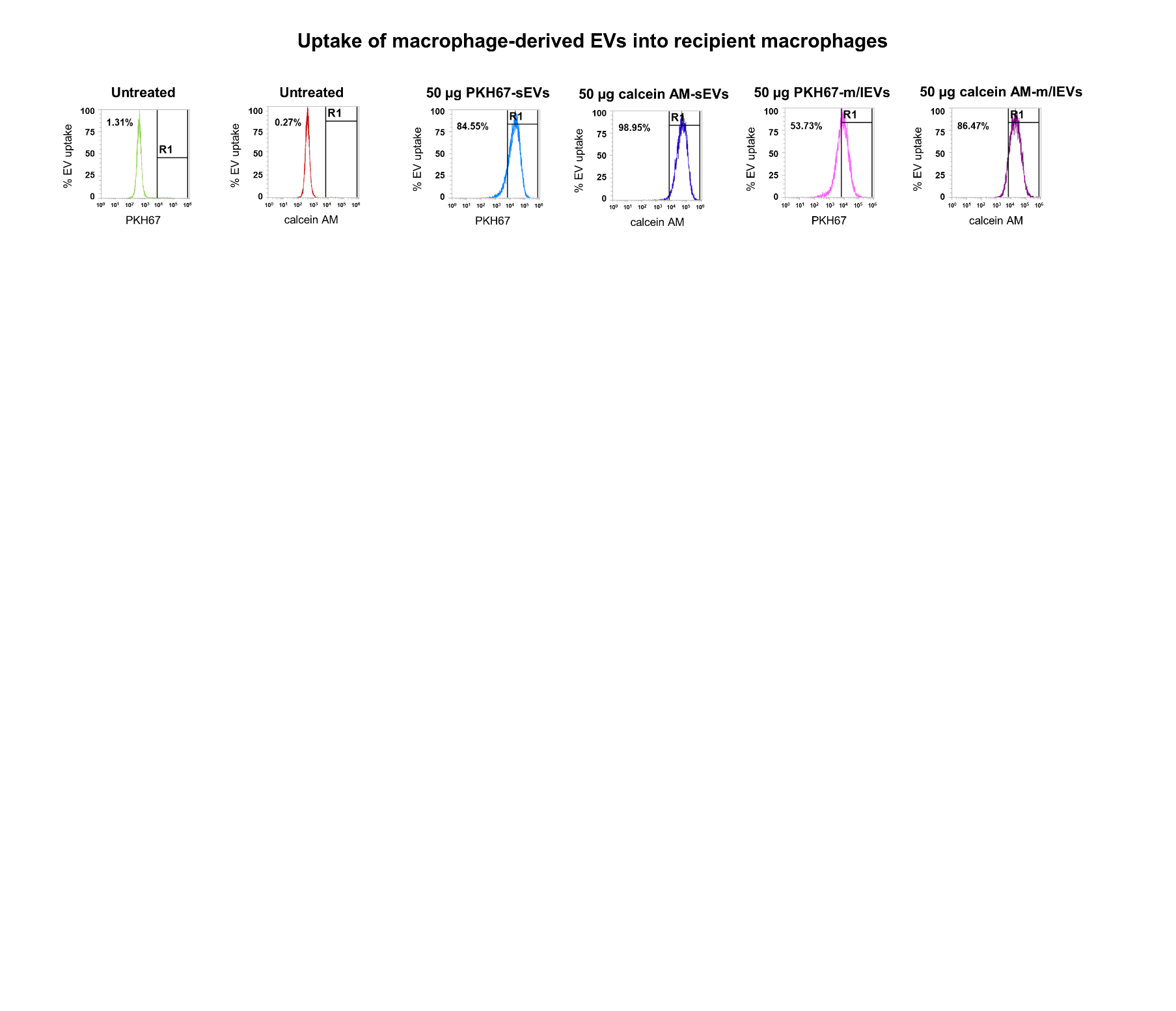

**Supplementary Figure 5. Representative histogram plots showing gating strategy for determining macrophage-derived EV uptake into recipient macrophages for figure 4 in main text.**

The recipient macrophages were treated with either PKH67- or calcein AM-labeled EVs at 5, 25, and 50 µg EV protein/well for 4 h. Data represent PKH67/calcein AM (+) cells noted by the shift past the solid black line after gating the autofluorescence of the untreated cells. The histogram plots are representative of triplicate samples.

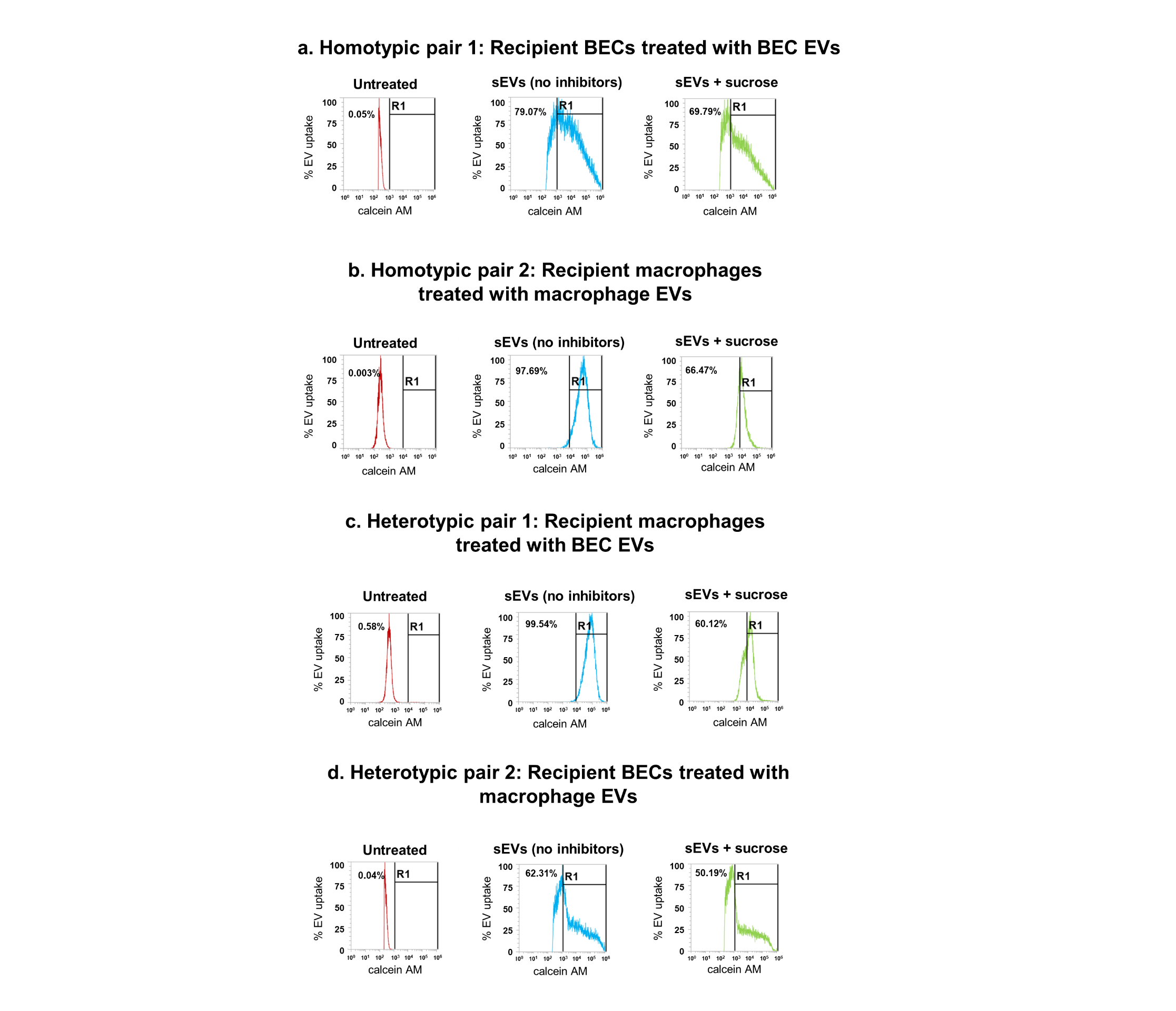

**Supplementary Figure 6. Representative histogram plots showing gating strategy for determining effect of endocytic inhibition on the cellular uptake of EVs (a, b, c, d) in figure 5 in main text.**

Recipient cells were pre-incubated for 0.5 h with endocytosis inhibitors and then were treated for 4 h m/lEVs and sEVs at 25 µg (recipient BECs) and 50 µg protein (recipient macrophages)/well in the presence of fresh endocytosis inhibitors. The % EV uptake and MFI values were determined using flow cytometry. Data represent calcein AM (+) cells noted by the shift past the solid black line after gating the autofluorescence of the untreated cells. The histogram plots are representative of triplicate samples.

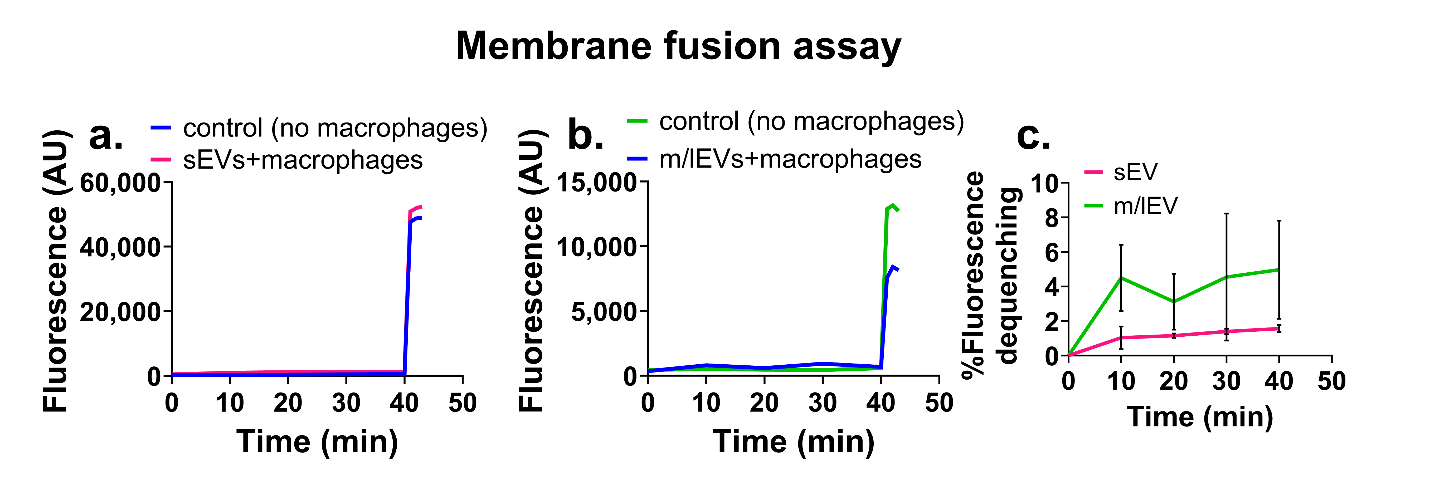

**Supplementary figure 7. Membrane fusion assay: Macrophage-derived EV uptake into recipient macrophages.**

Ten µg of R18-labeled EVs were added to MES buffer in a thermostated spectrofluorometer FluoroMax-4 and fluorescence was measured continuously at 560-nm excitation and 590-nm emission wavelengths (slit 1.5 nm). Post 10 min of equilibration, unlabeled macrophages were added to R18-EVs, and fluorescence was measured for a further 30 min. The fusion reaction was stopped by the addition of 0.3% Triton X-100 + 60 mM octylglucoside (**a, b**). R18 labeled sEVs or m/lEVs were left untreated (control) or mixed with unlabeled 1x10^6^ macrophages (sEV/m/lEV+cells) (**c**). Fluorescence dequenching (**FD**) values were calculated where %FD = ((F - F_i_ )/(F_max_ - F_i_)) x 100. Data is presented as mean±SD (n=3).
